# O-mannosylation and protein maturation check-points represent therapeutic opportunities in BRAF fusion protein oncogenesis

**DOI:** 10.1101/2025.08.18.670743

**Authors:** Sean A. Misek, Anna Borgenvik, Daniel Christen, Gloria Kyrila, Alexander Zhang, Sarah Reel, Michelle Boisvert, Kelly Cai, Kevin Zhou, Elizabeth M. Gonzales, Amy Goodale, Lorena Lazo de la Vega, Timothy J. Ragnoni, Zohra Kalani, Nicole Persky, Tanaz Abid, Esteban Miglietta, Sergey Vakhrushev, Michael Stumpe, Jacquelyn Jones, Seth Malinowski, Lobna Elsadek, Julie Parikh, Alexandra Larisa Condurat, Aaron Fultineer, Seung Hyun Choi, Merve Ozdemir, Zachary Eisenbies, Joohee Lee, Dana Novikov, Sher Bahadur, April Apfelbaum, Jenna Robinson, Kira Tang, Antonio Maldera, Daniel Bondeson, Jason Kwon, Mounica Vallurupalli, Hyesung Jeon, Sangita Pal, Todd Golub, William C. Hahn, Eric Fischer, Jesse S. Boehm, Jörn Dengjel, Henrik Clausen, Nada Jabado, Till Milde, Anne Carpenter, Beth A. Cimini, Keith Ligon, Katherine Janeway, Michael Eck, David Root, David T.W. Jones, Timothy N. Phoenix, Rameen Beroukhim, Hiren J. Joshi, Tilman Brummer, Adnan Halim, Pratiti Bandopadhayay

**Affiliations:** Department of Cancer Biology, Dana-Farber Cancer Institute, Boston, Massachusetts, USA; Department of Medical Oncology, Dana-Farber Cancer Institute, Boston, Massachusetts, USA; Broad Institute of MIT and Harvard, Cambridge, MA, USA; Department of Medicine, Harvard Medical School, Boston, MA, USA; Department of Pediatrics, Harvard Medical School, Boston, MA, USA; Dana-Farber/ Boston Children’s Cancer and Blood Disorders Center, Boston, MA, USA; Institute of Molecular Medicine, ZBMZ, Faculty of Medicine, University of Freiburg, Freiburg, Germany; Faculty of Biology, University of Freiburg, Freiburg, Germany; German Cancer Consortium (DKTK), Partner Site Freiburg and German Cancer Research Center (DKFZ), Heidelberg, Germany; Department of Cellular and Molecular Medicine, University of Copenhagen, Denmark; Cincinnati Children’s Hospital Medical Center, Cincinnati, Ohio, USA; Department of Biology, University of Fribourg, Fribourg, Switzerland; Department of Pathology, Dana-Farber Cancer Institute, Boston, Massachusetts, USA; Department of Biological Chemistry and Molecular Pharmacology, Harvard Medical School, Boston, MA, USA; Koch Institute for Integrative Cancer Research, Massachusetts Institute of Technology, Cambridge, MA, USA; McGill University, Quebec, Canada; University Hospital Jena, Department of Pediatrics and Adolescent Medicine, Friedrich Schiller University Jena, Jena, Germany; Comprehensive Cancer Center Central Germany (CCCG), Jena, Germany; Hopp Children’s Cancer Center Heidelberg (KiTZ), Heidelberg, Germany; Clinical Cooperation Unit Pediatric Oncology, German Cancer Research Center Heidelberg (DKFZ), Heidelberg, Germany; Division of Pediatric Glioma Research, Hopp Children’s Cancer Center Heidelberg (KiTZ), Heidelberg, Germany; National Center for Tumor Diseases (NCT), NCT Heidelberg, a partnership between DKFZ and Heidelberg University Hospital, Germany; German Cancer Research Center (DKFZ), Heidelberg, Germany; Division of Pharmaceutical Sciences, College of Pharmacy, University of Cincinnati, Cincinnati, OH, USA; Comprehensive Cancer Center Freiburg (CCCF), Medical Center, University of Freiburg, Faculty of Medicine, University of Freiburg, Freiburg, Germany; Center for Biological Signaling Studies BIOSS, University of Freiburg, Freiburg, Germany

## Abstract

Fusions between protein-coding genes are common oncogenic drivers across cancers, typically pairing a proto-oncogene with partner that does not independently drive cancer. In all therapeutically actionable fusions, the proto-oncogene is the drug target, the contributions to oncogenicity of the fusion partner have largely been ignored. We studied the role of BRAF fusion partners and found that they are necessary for transformation. In the setting of KIAA1549::BRAF, the most common fusion protein across brain tumors, we found that KIAA1549 is necessary for the oncogenicity of KIAA1549::BRAF and engenders a striking and specific dependency on the protein O-mannosyltransferase complex (POMT1/2). Specifically, we show that genetic silencing or pharmacologic inhibition of the protein O-mannosyltransferase complex (POMT1/2) reverses fusion-induced transformation, thereby representing a novel and MAPK independent therapeutic target. Furthermore, POMT1/2 is required to glycosylate and enable maturation of the K::B fusion protein. These findings represent a proof-of-concept for targeting the partners in oncogenic fusions as a potential cancer therapeutic strategy.

## Introduction

Since the introduction of imatinib as an inhibitor of BCR-ABL1 in patients with chronic myelogenous leukemia^1^, oncogenic fusion proteins have served as targets for the development of small molecule therapeutics. Typically, oncogenic fusions pair a single proto-oncogene (e.g. *ALK, NTRK, ROS1*, and *RET*) with any of a number of partner loci that do not encode known proto-oncogenes but nevertheless contribute to the amino acid sequence of the resulting protein^2^. An example is the proto-oncogene *BRAF*, which is fused in pediatric gliomas to one of multiple partners including *KIAA1549*^3–5^, *FAM131B*^6^, *RNF130*^7^, and *GNAI1*^7^. The constancy of the proto-oncogene and variability in its partner have directed most attention to understanding the role of the proto-oncogene component, with much less attention to the component encoded by the fusion partner. For example, *BRAF* fusions are thought to activate BRAF by truncating and removing autoinhibitory domains on its N-terminus, with reports suggesting that the fusion partners do not contribute to the oncogenicity of the protein product^8,9^.

*KIAA1549*::*BRAF* (K::B) rearrangements are particularly interesting as the sole and pathognomonic genetic alteration underlying most pilocytic astrocytomas (PAs) — the most common brain tumor in children^10^. K::B is challenging to therapeutically target since it retains a wildtype BRAF kinase domain, thus precluding the use of BRAF inhibitors developed to target BRAF^V600E^ in its monomeric state^11,12^. Recent clinical trials have evaluated on-target MAPK pathway inhibition in PAs and other pediatric low-grade gliomas (pLGGs) using either RAF or MEK inhibitors^13,14^. The type II BRAF inhibitor tovorafenib is approved for pLGG, but primary resistance, dose-limiting toxicity, and re-growth upon inhibitor withdrawal limit the utility of on-target MAPK inhibition in many patients^13,14^.

In this work, we assess whether proto-oncogene fusion partners are truly dispensable for oncogenicity, focusing on both common and rare *BRAF* fusion partners. We demonstrate that specific structural and functional elements of the *BRAF* fusion partners are necessary for transformation, in many cases by enhancing BRAF membrane localization. We also leverage genome-scale anti-transformation screens to demonstrate that *BRAF* fusion partners engender fusion-specific, therapeutically tractable, genetic dependencies. Most notably, we demonstrate that a necessary step for K::B oncogenicity is glycosylation of its KIAA1549 component by the protein O-mannosyltransferase 1 and 2 complex (POMT1/2), thereby suggesting this complex as a target for therapeutic development. This work highlights an underexplored approach to inhibit oncogenic fusions across pediatric and adult cancers.

## Results

### The role of BRAF fusion partners in pediatric low-grade gliomagenesis

Direct therapeutic targeting of oncogenic fusions has relied almost exclusively on inhibitors of the proto-oncogene component of the fusion. We performed an analysis of FDA-approved compounds for fusion-driven cancers to determine whether these target the proto-oncogene or fusion partner (**Fig. 1A**) and identified 21 FDA-approved compounds (**Supplemental Table 1**). In every case, the inhibitor targets the proto-oncogene product and not its fusion partner. This suggests that strategies to target the fusion partners remain largely underexplored.

**Figure 1.**
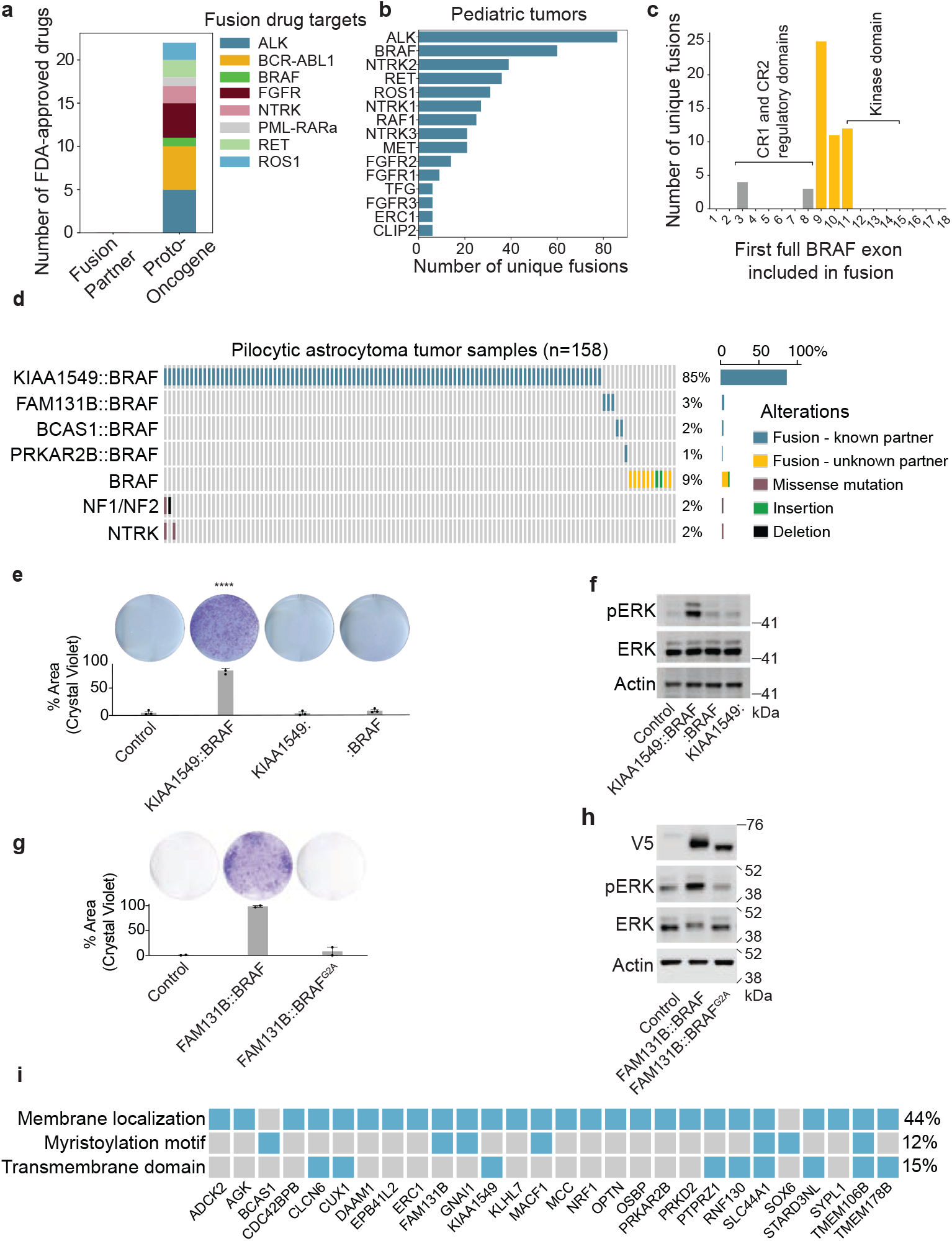
The role of BRAF fusion partners in pediatric low-grade gliomagenesis. **(A)** Bar graph of the number of FDA-approved drugs directly targeting either the fusion proto-oncogene or fusion partner. **(B)** Analysis of unique fusion species in tumors from 4,329 pediatric patients. Bars represent the number of unique fusion species identified (x-axis) for each fusion gene (y-axis). **(C)** Analysis of intronic breakpoint loci for 55 unique BRAF fusion species identified in an analysis of 4,329 tumors from pediatric patients. x-axis depicts the first complete exon downstream of the fusion breakpoint. y-axis depicts the total number of unique fusion species with a breakpoint in the given intron. **(D)** Genomic characterization of pediatric lowgrade glioma tumors. Recurrent driver alterations identified in a cohort of 158 Pilocytic Astrocytomas that have undergone targeted clinical sequencing. Driver alterations include structural variants (Fusions with a known partner and rearrangements with an unknown partner), insertions/deletions, and single nucleotide variants (SNVs). Columns depict individual tumor samples, and rows depict driver alterations. The fraction of tumors in the cohort with a given driver alteration are displayed as percentages. **(E)** Representative images and quantification of three independent colony formation assays of h9-NSCs expressing Luciferase (control), KIAA1549::BRAF, KIAA1549: (exons 1-15), and :BRAF (exons 9-18), and cultured in the absence of exogenous EGF and bFGF supplementation. Quantification (below) of the percentage of plate area with cells. Error bars represent Standard Error of the Mean (SEM) across biological replicates, one-way ANOVA p < 0.0001***. **(F)** Representative immunoblot showing protein levels of phosphorylated and total ERK in h9-hNSCs expressing the same constructs as in (d). Lysates were collected 24 hours after withdrawal of exogenous EGF and bFGF supplementation. **(G)** Representative images and quantification of two independent colony formation assays of h9-hNSCs expressing Luciferase (control), FAM131B::BRAF, and FAM131B::BRAF^G2A^ and cultured in the absence of exogenous EGF and bFGF supplementation. **(H)** Representative V5, pERK, ERK, and Actin immunoblots from h9-NSCs expressing the same constructs as in (f). Luciferase (control), FAM131B::BRAF, and FAM131B::BRAF^G2A^ constructs all have. C-terminal V5 tag. Data are from two independent replicates. Quantification of immunoblots in (**Supplemental Fig. 1B**). **(I)** Analysis of n = 28 (of a total of 59) BRAF fusion partners identified in an analysis of 4,329 pediatric cancers. Heatmap displays those with evidence for membrane localization (on Uniprot^114^), an N-terminal myristoylation motif, or one or more transmembrane domains. A complete set of BRAF fusion partners is described in (**Supplemental Table 3)**.

Reasoning that fusion partners may provide additional targets for fusion-driven cancers, we were interested in understanding the heterogeneity of fusion partners for the most common proto-oncogenes across adult cancers, and all those involving kinases in pediatric tumors (**Fig. 1B and Supplemental Fig. 1A**). Across pediatric tumors with kinase fusions, we identify 770 unique fusion species, with ALK having the most (n = 86) followed by BRAF (n = 60). Interestingly, in a second dataset of predominantly adult tumors (FusionGDB^15^), BRAF also had the second greatest number of unique fusions, with 37 identified across 8101 tumors samples with at least one gene fusion. This heterogeneity in BRAF fusion partners is consistent with previous analyses^16^. Given the frequency of BRAF fusions across pediatric and adult cancers, and the large number of fusion variants, we sought to evaluate whether fusion partners may present unexplored therapeutic potential, using BRAF as an example.

BRAF fusions have largely been thought to induce oncogenesis through truncation and removal of critical negative regulatory domains on the N-terminus of BRAF, thereby rendering the kinase domain constitutively active^8,17^ BRAF is composed of N-terminal regulatory domains (CR1 and CR2) and a C-terminal kinase domain (CR3). CR1 and CR2 act as autoinhibitory regions and are essential for recruitment of BRAF to the plasma membrane^18^. In every BRAF fusion present in our pediatric cohort, the rearrangement disrupts at least CR1, resulting in deletion of these N-terminal auto-inhibitory domains (**Fig. 1C**). However, the loss of these residues also necessarily includes loss of the regions in BRAF that mediate physical interactions with RAS and the localization of BRAF at the plasma membrane, which are necessary for placing it in position for activation^11,19^. The loss of these BRAF domains suggests a role for the BRAF fusion partners in localizing truncated BRAF to the plasma membrane to enable its activation. Consistent with this hypothesis, every BRAF fusion adds non-BRAF amino acids to the truncated BRAF kinase. This is in contrast to enhancer-hijacking fusions, in which upstream, non-coding regulatory elements of the 5’ gene partner are hijacked to drive overexpression of the 3’ effector^7,20,21^. We therefore set out to understand the contributions of BRAF fusion partners, with the hypothesis that they contribute to the cellular localization of the fusion oncogene.

BRAF fusions are pathognomonic for pilocytic astrocytoma (PA), the most common subtype of pediatric low-grade glioma (pLGG)^10^. BRAF fusions are typically the only genetic alteration across a PA’s genome. The vast majority (>90%) fuse C-terminal BRAF to N-terminal KIAA1549 (**Fig. 1D, Supplemental Table 2**)^10^. We therefore decided to focus our studies on the role of KIAA1549.

A major challenge to the study of PAs is a lack of patient-derived model systems; PAs cannot be propagated *in vitro* or as patient-derived xenografts using standard methods. SV40-immortalized models have been generated^22,23^, but these do not retain dependency on K::B. As an alternative strategy to generate oncogene-relevant model systems, we engineered both murine and human neural stem cell (NSC)-derived PA models by exogenously expressing K::B. We started with h9 embryonic stem cell-derived human neural stem cells (h9-NSCs) that we engineered to express IKZF3 degron-tagged K::B, BRAF^V600E^, and luciferase controls (**Extended Fig. 1A and B**). In this system, pomalidomide induces the interaction between the IKZF3 tag and the CRBN-CRL4 E3 ligase, resulting in rapid titratable degradation of tagged proteins^24^ (**Extended Fig. 1C**). Expression of K::B or BRAF^V600E^ using this system is sufficient to transform hNSCs as measured by *in vitro* growth in the absence of exogenous EGF and bFGF supplementation (**Extended Fig. 1D**). In contrast, the luciferase controls remained dependent on exogenous EGF and bFGF (**Extended Fig. 1D**). Growth of oncogene-transformed h9-NSCs was blocked by pomalidomide-induced degradation of K::B or BRAF^V600E^, confirming specificity of this phenotype to oncogene expression (**Extended Fig. 1E**). Similarly, degradation of K::B or BRAF^V600E^ prevented induction of ERK phosphorylation and proliferation (**Extended Fig. 1F and G**). These data indicate that expression of either form of activated BRAF is sufficient for transformation of human neural stem cells.

We then used these models to determine whether KIAA1549 is necessary for K::B-induced transformation, using proliferation in the absence of exogenous EGF/bFGF as the transformation phenotype. To this end, we expressed truncated variants of K::B with only the KIAA1549 portion of the fusion (K:) or the BRAF portion of the fusion (:B) and compared their transforming potential to full length K::B or the luciferase control. Expression of full-length K::B, but not the K: or :B truncation mutants, significantly induced two-dimensional colony formation (**Fig. 1E**; p < 0.0001) and MAPK pathway activation (**Fig. 1F**). This observation indicates that KIAA1549 is indispensable for K::B activation.

We suspect that this is true for all BRAF fusion partners in oncogenic fusions. We performed a meta-analysis of kinase-driven fusions in pediatric solid tumors and cataloged characteristics of fusions described in 531 papers published between 2013 and 2022 (**Supplemental Table 3**). This analysis identified 31 unique protein coding BRAF fusion partners in pediatric brain tumors including PAs. Among these fusion partners, the partner with the smallest contribution to the fusion protein was *FAM131B*, which appends as few as nine FAM131B residues to the N-terminus of truncated BRAF^6^. Strikingly, adding these nine amino acids to truncated BRAF was sufficient to transform h9-NSCs (**Fig. 1G**) and activate the MAPK pathway (**Fig. 1H** and **Supplemental Fig. 1B**; p < 0.0001).

Intriguingly, the first six residues of the FAM131B::BRAF fusion encode a canonical MGXXXS myristoylation motif^25,26^. Myristoylation is a post-translational modification that adds a lipid to the N-terminus of a protein, thereby facilitating attachment of the protein to membranes^27^. This observation led us to suspect that myristoylation of the fusion protein mediates oncogenesis by attaching it to membranes, resulting in increased local concentrations and the dimerization required for aberrant activation^17,28–31^. We therefore tested the necessity of myristoylation to the oncogenicity of FAM131B::BRAF by mutating the second FAM131B amino acid, glycine, to alanine, abolishing the myristoylation site. This completely prevented FAM131B::BRAF-induced transformation (22-fold decrease relative to FAM131B::BRAF) (**Fig. 1G**) and MAPK pathway activation (4.17-fold change, p < 0.0001) (**Fig. 1H** and **Supplemental Fig. 1B**). Since myristoylation tethers proteins to intracellular membranes, we also assessed subcellular localization of FAM131B::BRAF and FAM131B::BRAF^G2A^ with immunofluorescence microscopy. FAM131B::BRAF predominantly localized to the plasma membrane (**Supplemental Fig. 1C**), whereas FAM131B::BRAF^G2A^ did not — consistent with the idea that myristoylation of FAM131B::BRAF enhances membrane localization.

We were interested in understanding whether additional BRAF fusion partners harbored domains that may similarly facilitate recruitment of truncated BRAF to the membrane, allowing activation of downstream MAPK signaling. We therefore performed a computational analysis of 59 BRAF fusion partners identified in pediatric cancer (**Supplemental Table 4**) to evaluate whether they harbor domains that might facilitate membrane tethering (**Fig. 1I**), including transmembrane domains and myristoylation sites. Across all 59 fusion partners, seven contained myristoylation sites and an additional nine, including KIAA1549, were predicted to contain at least one transmembrane domain for at least one isoform. A variety of lipid modifications in addition to myristoylation can also localize proteins to membranes, and a UniProt subcellular localization survey of all 59 BRAF fusion partners identified an additional 10 that localized to a membrane, for 26 (44%) overall (**Fig. 1I**).

### Anti-transformation screens identify additional dependencies in KIAA1549::BRAF-vs BRAF^V600E^-transformed cells

Given that KIAA1549 is necessary for oncogenicity of the K::B fusion, we hypothesized that additional genes and pathways beyond the canonical MAPK pathway may also be required. We also reasoned that these would not be shared by activating point mutations like BRAF^V600E^. To test this, we performed genome-scale loss of function CRISPR/Cas9 screens in embryonic murine neural stem cells (mNSCs) expressing human K::B (the 15::9 fusion breakpoint variant), human BRAF^V600E^, or luciferase (**Fig. 2A** and **Extended Fig. 2A**). We chose mNSCs because we had previously optimized conditions for growing mNSCs sufficiently to conduct genome-scale screens^43,44^. As with hNSCs, expression of K::B or BRAF^V600E^ was sufficient to transform mNSCs as measured by anchorage-independent *in vitro* growth as neurospheres in the absence of exogenous EGF and bFGF supplementation (p<0.001) (**Extended Fig. 2B-D**), and by glioma formation *in vivo* when injected as intracranial allografts^45^, further indicating that both K::B fusions and BRAF^V600E^ are sufficient for transformation. In lieu of a standard synthetic lethal screen, we designed this screen to also assess cofactors that contribute to growth factor independence, the transformation phenotype of K::B and BRAF^V600E^. Specifically, we grew the K::B- and BRAF^V600E^-expressing cells in the absence of the growth factors EGF and bFGF, rendering these cell lines oncogene-dependent. Both synthetic lethalities and transformation cofactors will appear as dependencies in this screen. The luciferase controls were grown in standard media containing EGF and bFGF (**Fig. 2A** and **Extended Fig. 2B-D**).

**Figure 2.**
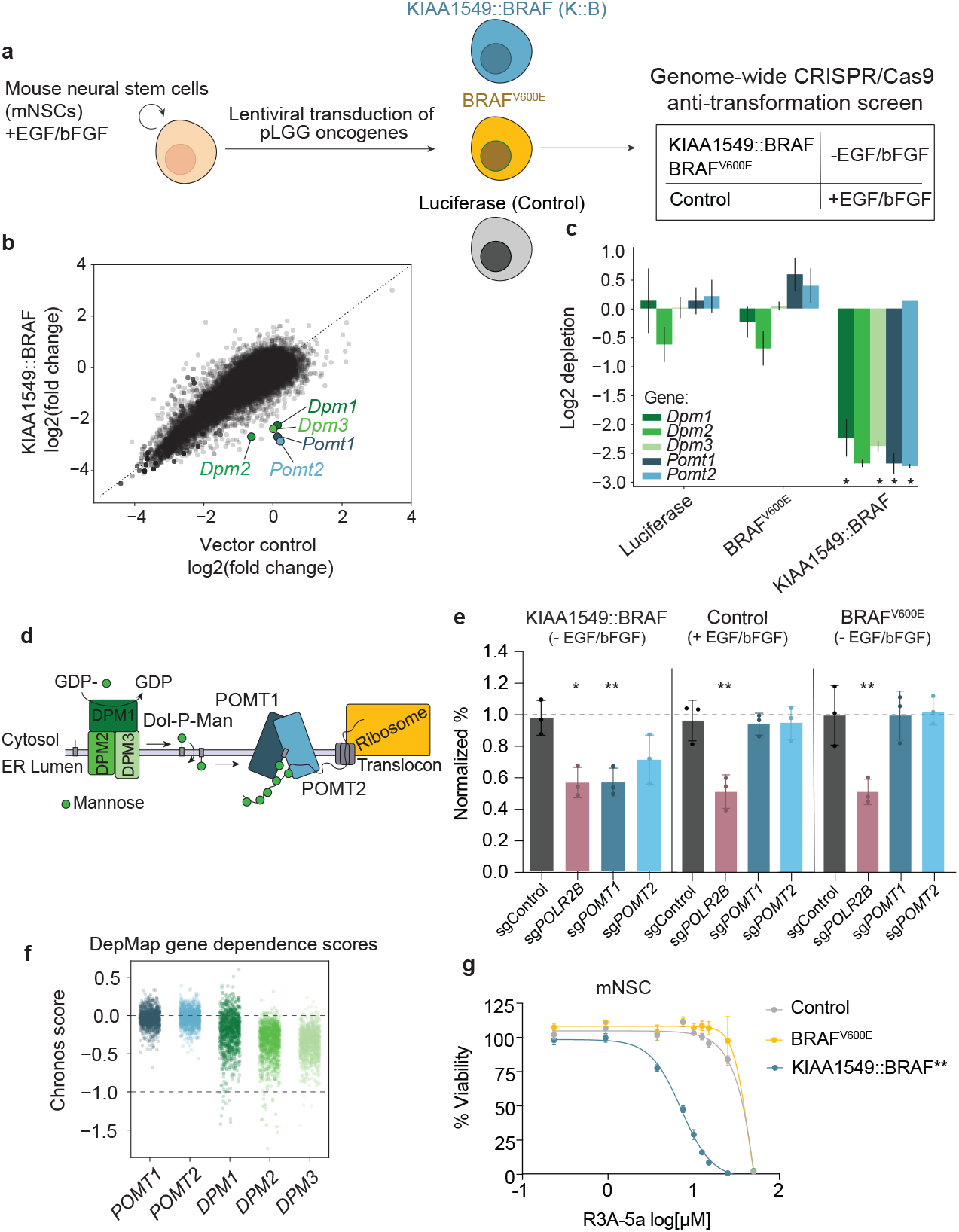
Anti-transformation screens identify additional dependencies in KIAA1549::BRAF- vs BRAFV600E-transformed cells. **(A)** Schematic depicting generation of isogenic cell models from embryonic (E14.5) mouse neural stem cells with ectopic expression of human KIAA1549::BRAF (15:9), BRAF^V600E^, and Luciferase (Control). Isogenic lines with expression of KIAA1549::BRAF and BRAF^V600E^ were cultured without EGF/bFGF, rendering them oncogene dependent. Luciferase-expressing cells were cultured with EGF/bFGF. These models were subsequently leveraged to perform genome-scale loss of function CRISPR/Cas9 screens (**Extended Fig. 2**). (**B**) Scatter plot showing log_2_ fold change (day 18 vs early timepoint) in gene-level CRISPR guide abundance scores. Screen from mNSCs expressing KIAA1549::BRAF (in the absence of EGF/bFGF) is represented on the y-axis. Screen from mNSCs expressing Luciferase (in the presence of EGF/bFGF) is represented on the x-axis. Each dot represents a single gene, selected genes are highlighted. Each screen was performed with three independent biological replicates. **(C)** Bar graph of log(fold change) gene-level depletion values for selected genes in the genome-scale CRISPR/Cas9 screen described in Fig. 2B. Each log(fold change) value is computed by comparing the log(fold change) at the final (day 18) timepoint vs the early (day 3) timepoint. * value depicts p value <0.05 as calculated by Student T-tests comparing log fold change in K::B expressing cells compared to luciferase or BRAF^V600E^ controls. **(D)** Graphical representation of POMT1/POMT2 specific O-Mannosylation in mammals. The dolichol phosphate mannose (Dol-P-Man) donor is synthesized from Dol-P and GDP-Man by the DPM1-3 complex. Dol-P-Man flips and is utilized by the POMT1/POMT2 heterodimer for co/post-translational modification of Ser/Thr residues with O-linked mannose within unstructured protein domains in the ER lumen. DPM1/2/3: dolichol-phosphate mannosyltransferase-1/2/3 of the Dol-P-Man complex; GDP/GTP: guanosine diphosphate/triphosphate; POMT1/2: protein O-mannosyltransferase 1 and 2 of the POMT complex. **(E)** Bar plot of CRISPR competition assay in h9-NSCs. Human neural stem cells were engineered to express KIAA1549::BRAF, BRAF^V600E^, or Luciferase and cultured as described in (a). Cells were transduced with either intergenic control guides (EGFP) or guides targeting ***POLR2B, POMT1, POMT2***, or the same intergenic control regions (mCherry). Three independent guide sequences were used for each condition. Cells were pooled 1:1 as described in (**Supplemental Table 5**). The fraction of mCherry- or EGFP-positive cells was assessed with flow cytometry three days post-transduction (initial timepoint) and 18 days post-transduction (final timepoint). The fraction of mCherry positive cells, normalized to the initial timepoint, is displayed on the y-axis. The sgRNA pair conditions are displayed on the x-axis. Statistical analysis was performed using one-way ANOVA with Dunnett’s multiple comparison test (vs sgControl). Statistically significant p-values (p < 0.05) are indicated with * on the plot. **(F)** Dot plot with Chronos^109^ gene dependency scores for ***POMT1, POMT2, DPM1, DPM2*** and, ***DPM3*** across 1150 cancer cell lines profiled in The Cancer Dependency Map (DepMap)^109^. A Chronos score of 0 or −1 is the median of all genes classified in the DepMap 24q2 release as non-essential or common essential, respectively. **(G)** Best-fit curve of mNSC isogenic models expressing Luciferase (control), KIAA1549::BRAF, and BRAF^V600E^ treated with increasing concentrations of the POMT complex inhibitor R3A-5a. Viability was measured with CellTiterGlo after 72 hours of treatment. Cell viability was normalized to each equivalent DMSO control. Viability was assessed using area under the curve (AUC) of each experimental replicate (n = 3), and then compared using one-way ANOVA with Sidak’s multiple comparisons test (luciferase vs KIAA1549::BRAF p = 0.0009^***^, luciferase vs BRAF^V600E^ p=0.7). AUC luciferase control = 220.8; KIAA1549::BRAF = 140.8; BRAFV600E = 233).

Strikingly, K::B expressing cells exhibited strong and selective dependencies that were not observed across the other mNSC models (BRAF^V600^E or luciferase), with dependencies on *Pomt1* and *Pomt2* standing out as the most significant (Pomt1 differential logfc = −2.82, *Pomt2* differential logfc = −3.07, Pomt1 p = 0.0006, *Pomt2* p = 0.0004) (**Fig. 2B and C**). Pomt1 and Pomt2 heterodimerize to form the POMT1/2 complex^35^. POMT1/2 catalyzes O-mannosylation, transferring mannose from the Dol-P-Man donor to Ser and Thr residues on acceptor protein substrates^41,47^ (**Fig. 2D**). In addition, the genetic dependency on *Pomt1/2* uncovered by our anti-transformation screens was further supported by highly selective K::B cell dependencies on all three members of the heterotrimeric DPM complex (*Dpm1, Dpm2*, and *Dpm3*) (**Fig. 2B and C**). *Pomt1, Pomt2, and Dpm1-3* all exhibited greater than 65% depletions (log2 change <-1.5; *Dpm1* p = 0.021, *Dpm2* p = 0.0024, *Dpm3* p = 0.0023) over the 18-day screen. Intriguingly, in unrelated work, we had previously showed that wild-type KIAA1549 is the most heavily O-mannosylated protein across the human proteome41 and that KIAA1549 O-mannosylation is mediated by POMT1/2^47,48^. Taken together, these findings indicate that K::B expressing NSCs harbor a striking dependency on Dol-P-Man donor availability, POMT1/2 activity, and O-mannosylation of the extracellular K::B domain

### Expression of POMT1 and POMT2 in cancer and normal brain

To assess whether O-mannosylation by the POMT1/2 complex might have a role in human pLGGs with K::B fusions, we analyzed expression of both POMT1/2 and DPM complex members across a panel of 87 human K::B-positive pLGGs profiled with bulk RNA-Seq. We detected POMT1, POMT2, and all three members of the DPM complex in every tumor profiled, with read counts of at least 8 per million reads (average across all genes = 67.3; range 11.9-261.1) (**Extended Fig. 3A/B**). Tumors from pLGG patients typically have high non-malignant cell infiltration^49^, confounding bulk RNA-seq analysis since many sequencing reads are derived from non-malignant cells. We therefore interrogated single cell RNA-seq data from 10 K::B-driven pLGGs comprising 60,131 cells (average per tumor: 6013 cells; range 912-16,633). Because these tumors have few genetic alterations, it can be difficult to distinguish between cancer and normal cells in these data. However, our prior analyses of single-cell RNA-seq data from K::B-driven pLGGs indicated that the vast majority of glia exhibited the fusion^49^. We therefore performed Leiden clustering and manual annotation of each cluster according to 10 prior gene sets. We detected nine different cell types (**Supplemental Fig. 2**). POMT1/2 and DPM complex genes were preferentially expressed across glia and neurons relative to immune infiltrates (myeloid cells and T-cells) and endothelial cells (p < 0.001) (**Supplemental Fig. 2**).

K::B-driven pLGGs predominantly arise in the cerebellum^10^, and the high expression of *POMT1* and *POMT2* seen in K::B driven pLGGs appears to be consistent with expression patterns in normal brain. We analyzed expression of *POMT1, POMT2, DPM1, DPM2*, and *DPM3* in bulk RNA-seq (GTEx^50^) data spanning 54 normal tissue types, including 13 brain regions. Both *POMT1* and *POMT2* exhibited preferential expression in the brain, with highest expression in the cerebellum (**Supplemental Fig. 3**). DPM complex genes were more uniformly distributed across tissue types (**Supplemental Fig. 3**), reflecting their pleiotropic functions beyond O-mannosylation^51–53^. We conclude that PAs with K::B fusions arise in brain locations that express high levels of POMT1/2 complex genes relative to other tissues, and that the PAs retain this expression.

### POMT1 and POMT2 are necessary for KIAA1549::BRAF-induced transformation

Next, we validated *Pomt1* and *Pomt2* as genetic dependencies across our panel of isogenic mNSCs and hNSCs. We used CRISPR/Cas9 to silence *Pomt1* or *Pomt2* in K::B-expressing or control mNSCs and assessed proliferation over 18 days. We included cells expressing guides against GFP as a non-targeting negative control. Loss of *Pomt1* or *Pomt2* significantly reduced proliferation of K::B-dependent mNSCs (p= 0.007 and, p= 0.02 respectively), but not luciferase-expressing controls (p=0.92 and p=0.95) (**Supplemental Fig. 4A**). CRISPR cut-site sequencing of K::B-expressing mNSCs that survived to day 18 provided further evidence of a POMT dependency: almost all reads (91% and 94%, respectively) had either no editing or in-frame indels at the targeted loci, indicating that these surviving K::B-expressing mNSCs retained functional *Pomt1/2*. In contrast, 74% and 79% of luciferase control mNSCs exhibited frameshift mutations at the *Pomt1* and *Pomt2* cut sites, respectively. Indeed, whereas the proportion of reads with frameshift mutations at the targeted loci increased over time in these control mNSCs, they decreased in K::B-expressing mNSCs, suggesting that K::B cells with frameshift mutations were selectively lost (**Supplemental Fig. 4B**). Similarly, CRISPR/Cas9 ablation of *POMT1* or *POMT2* was sufficient to impede growth of K::B-expressing hNSCs relative to the non-targeting vector controls (**Fig. 2E**, p = 0.0403) to a similar degree as ablation of the pan-essential gene *POLR2B* (**Supplemental Table 5 and Supplemental Fig. 4C**, luciferase p = 0.002, BRAF^V600E^ p = 0.001, K::B p = 0.034). We observed no differences following suppression of either *POMT1* or *POMT2* in control hNSCs engineered to express BRAF^V600E^ or luciferase (**Fig. 2E**, luciferase p = 0.98, BRAF^V600E^ p = 0.98).

Indeed, dependence on POMT1/2 complex members is extremely specific to K::B-dependent cells. None of the 1150 cell lines profiled with genome-scale CRISPR screens in the Cancer Dependency Map^54^ contain a K::B fusion, and none exhibited a strong dependency on either *POMT1* or *POMT2* (Chronos^55^ score <-1) (**Fig. 2F**). A subset of cell lines (n = 22/1150, 2%) did exhibit dependency on one or more members of the DPM complex (**Fig. 2F**), potentially due to pleiotropic functions of Dol-P-Man that are unrelated to the POMT1/2 complex^51–53^. We conclude that the POMT1/2 complex is *specifically* essential for the proliferation of K::B-dependent NSCs.

This dependence on POMT1/2 appears to relate to its enzymatic activity. We treated our isogenic mNSC models with R3A-5a, a competitive inhibitor of the active site of the yeast ortholog of POMT1/2, Pmt1/2^56,57^. R3A-5a impeded growth of mNSCs expressing K::B, but not BRAF^V600E^ or the luciferase vector control (AUC K::B: 141, BRAF^V600E^: 233, Luciferase control: 221; p = 0.0009 and p = 0.7) **Fig. 2G** and **Supplemental Fig. 4D**).

### The POMT1/2 complex O-mannosylates K::B in the ER

Our finding that K::B expressing cells are exquisitely dependent on the POMT1/2 complex led us to hypothesize that, like wild-type KIAA1549, the K::B fusion is also heavily glycosylated and that this is necessary for its oncogenic function. We first confirmed that K::B is glycosylated by performing enzymatic digestion of the fusion protein. Treatment of lysates from K::B-expressing cells with PNGase F, which removes all N-linked glycans, induced an apparent mass shift of K::B of at least 20 kDa in immunoblots (**Supplemental Fig. 5A**). This magnitude of difference in the molecular mass corresponds to approximately 8-10 occupied N-glycosylation sites, consistent with the 10 potential N-glycosylation sites N-terminal to the TM domain of KIAA1549 (residues 1299-1319).

To further confirm that K::B undergoes O-mannosylation in our mNSC model, we combined lectin enrichment and differential O-glycoproteomics (**Fig. 3A**)^47,48^. Total cell extracts of control and K::B expressing mNSC cells were differentially labeled with diethyl stable isotopes (**Supplemental Fig. 5B**) and enriched for O-Man glycosylated peptides for identification and relative quantification by mass spectrometry (**Fig. 3A**). We identified 28 O-Man glycosylated proteins in mNSCs, including O-Man glycosylation of endogenous mouse Kiaa1549 and the stably expressed human K::B fusion protein (**Fig. 3B-D**). O-Man glycopeptides derived from endogenous Kiaa1549 were present at ~1:1 heavy-to-light ratios. These findings confirm that wild-type mouse Kiaa1549 substrate is expressed and acquires O-Man glycans in comparable amounts in control and K::B-expressing mNSCs, consistent with our previous findings^47,48^. In contrast, we identified human K::B O-Man glycopeptides exclusively with heavy stable isotope labels. This showed that human K::B is expressed and O-Man glycosylated only in the human K::B-expressing mNSC cells (**Fig. 3B**). We identified 32 O-Man glycosites on the K::B fusion protein, all of which were N-terminal to the TM domain (K::B residues 1299-1319). Collectively, these results demonstrate that K::B is expressed as a type I transmembrane protein in mNSCs and that the N-terminal domain acquires O-Man glycans in the ER lumen, where the Pomt1/2 complex is active^58^. Together, our data confirm that the K::B fusion protein is heavily mannosylated by the POMT complex.

**Figure 3.**
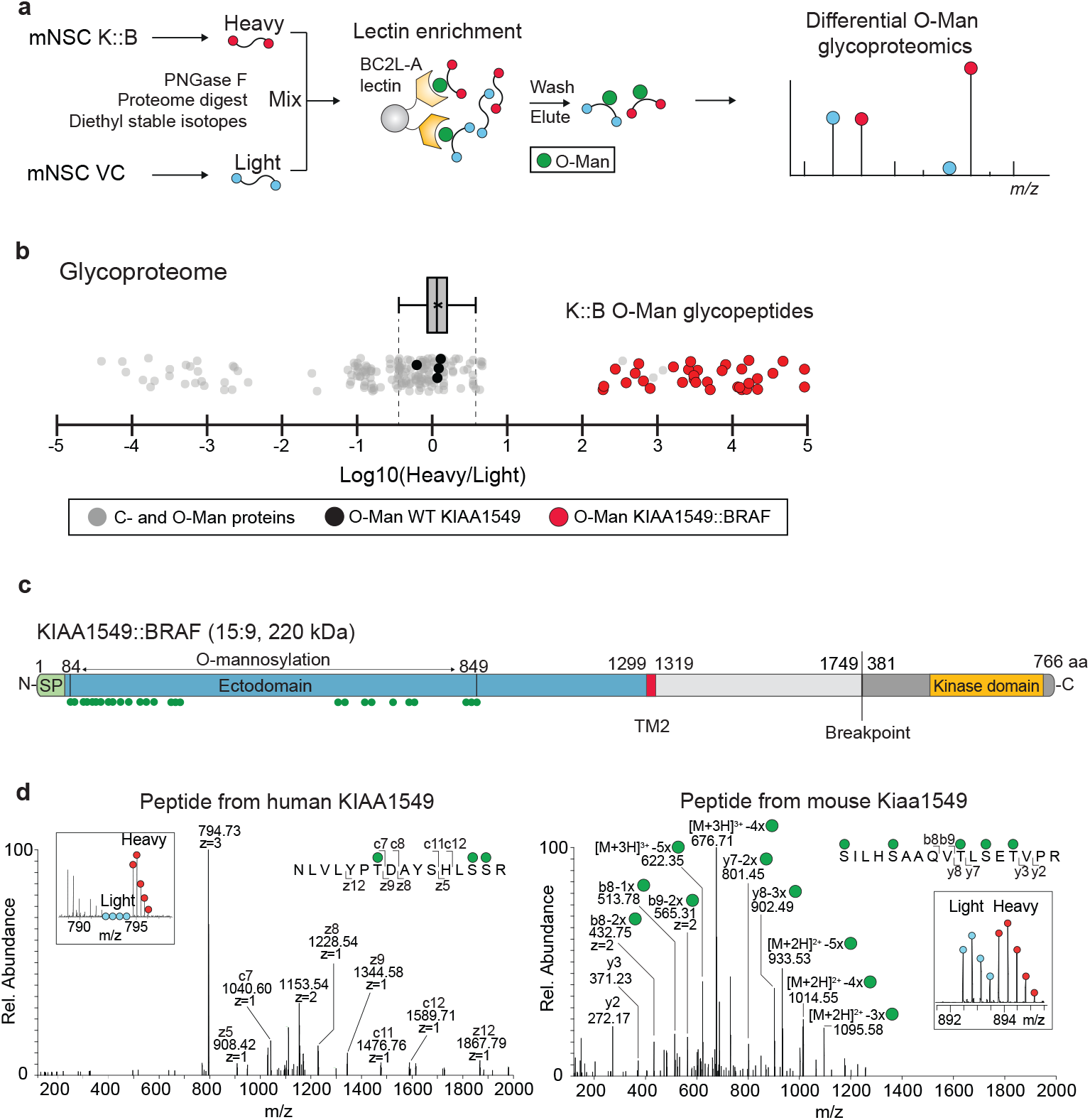
The POMT1/2 complex O-mannosylates K::B in the ER. **(A)** Workflow for quantitative O-Man glycoproteomics of stable isotope labelled total cell digests enriched for O-Man glycopeptides by BC2L-A lectin chromatography. **(B)** The relative quantification of the total proteomes is shown as boxplot with whiskers for which the inter quartile range calculations were used to determine the boundaries (dashed lines) for peptide spectral matches (PSMs) derived from Luciferase- or human KIAA1549::BRAF-expressing mNSCs. Dot-plot shows heavy-to-light (H/L) ratios of individual O-Man PSMs from KIAA1549:BRAF (red), endogenous mouse KIAA1549 (black) and other C- and O-Man glycoproteins (grey), demonstrating that O-Man of endogenous mouse KIAA1549 is equally abundant while O-Man glycopeptides of human KIAA1549::BRAF are exclusively derived from the mNSCs engineered to express human KIAA1549::BRAF. **(C)** Schematic illustration of human KIAA1549::BRAF showing the mapped O-Man glycans (green spheres) to residues 84-849 of the N-terminal domain. **(D)** Representative PSMs used for identification and site mapping of O-Man glycans on human KIAA1549::BRAF (left) and endogenous mouse KIAA1549 (right). Insert shows precursor isotope envelopes for heavy (red) and light (light blue) labelled glycopeptides selected for MS2 fragmentation.

### K::B, unlike wild-type BRAF, traffics through the secretory pathway

Wildtype KIAA1549 is predicted to contain transmembrane domains, which are retained in K::B, and we assessed their necessity for oncogenic activity. Initial modeling suggested that KIAA1549 has an N-terminal signal peptide (residues 1-61) and two transmembrane domains (residues 998-1018 and 1299-1319), but this is inconsistent with oncogenic activity of K::B since BRAF would not be exposed to the cytoplasm and its downstream effector MEK. We therefore characterized the orientation of KIAA1549 using three different approaches. First, we inferred protein topology with DeepTMHMM^32^ to identify intracellular, extracellular, and transmembrane domains (**Fig. 4B**). This analysis identified N-terminal residues 1-61 of KIAA1549 as a signal peptide and predicted that the remaining residues before residue 1299 are extracellular. All residues after 1319 are predicted to be intracellular, suggesting that residues 1299-1319 reside in the membrane. Second, we computed local hydrophobicity across KIAA1549 as previously described^33^ and identified this locus as the only region with enrichment of hydrophobic residues (**Fig. 4B**). Third, we analyzed the predicted structure of KIAA1549 with AlphaFold^55,56^ and identified only a single transmembrane helix matching the topology prediction (**Extended Fig. 4**). These analyses suggest that only residues 1299-1319, which coincide with the second of the two previously predicted transmembrane domains, truly cross the membrane. The other putative transmembrane helix (AA 998-1018) is predicted by AlphaFold to be part of a larger structured domain with high homology to SEA (Sea urchin sperm protein, Enterokinase, Agrin) domains^36^ in the extracellular-facing side of the protein (**Extended Fig. 4**).

**Figure 4.**
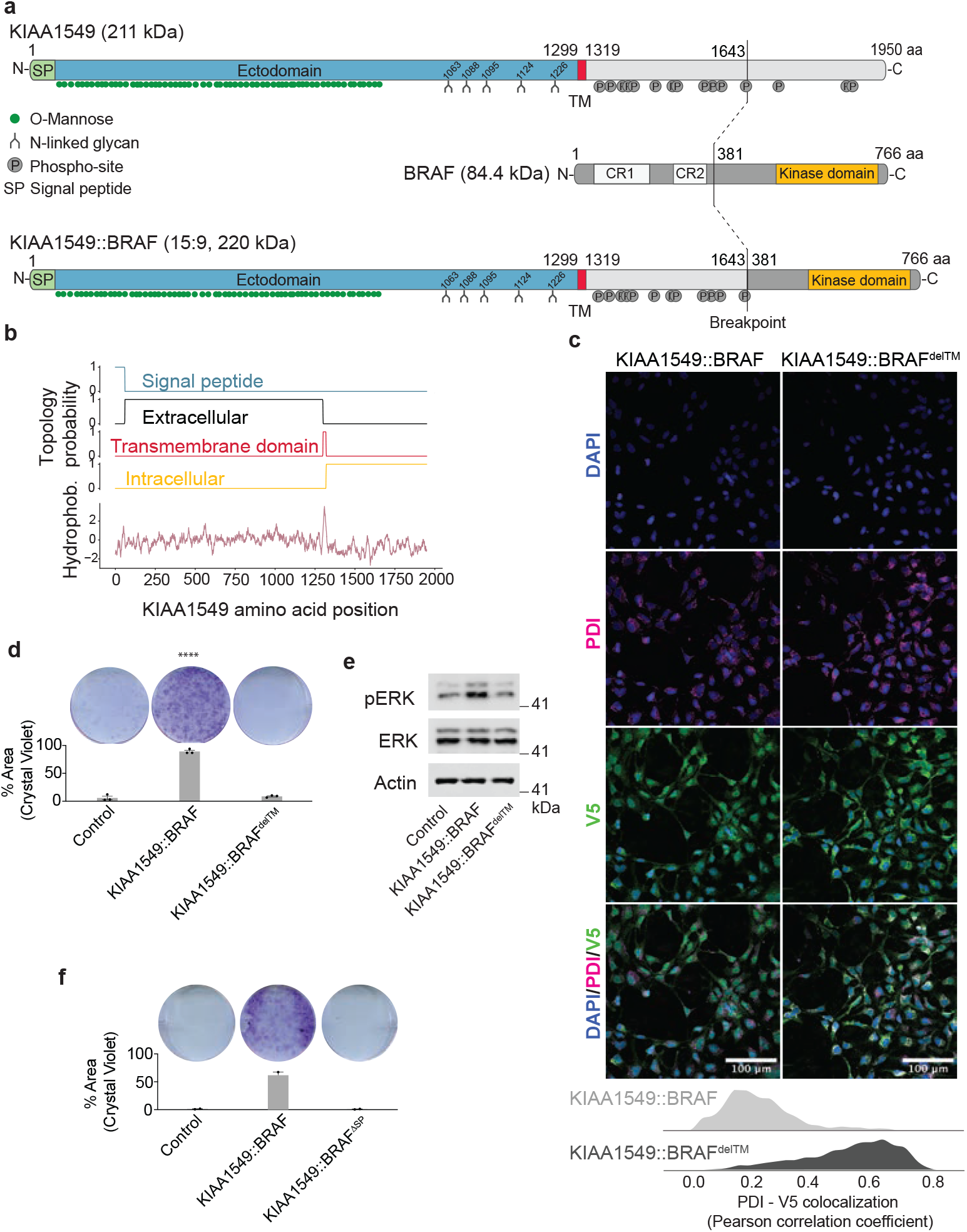
K::B, unlike wild-type BRAF, traffics through the secretory pathway. **(A)** Schematic showing wildtype human KIAA1549 (top panel), wildtype human BRAF (middle panel), and KIAA1549::BRAF (15::9, lower panel), annotated to show the presence of glycosylation or phosphorylation modifications. Approximate location of n = 15 phosphorylation sites are annotated according to Uniprot^114^. N-glycosylation (n = 5) and O-glycosylation (n = 64) sites are annotated as forks or green circles, respectively. The KIAA1549 ectodomain (residues 62-1299) is annotated as all residues preceding the KIAA1549 transmembrane domain (residues 1299-1319) excluding the ER signal peptide (residues 1-61). SP: signal peptide; TM: transmembrane domain; CR1: conserved region 1 (RAS-binding domain and cysteine-rich domain); CR2: conserved region 2 (autoinhibitory hinge region). **(B)** Transmembrane topology of wildtype KIAA1549 was predicted with DeepTMHMM^24,102^ (top). Local hydrophobicity^24,108^ (bottom) was predicted using the approach described in Kyte and Doolittle. Both analyses identify a single transmembrane domain (residues 1299-1319) in wildtype KIAA1549, that is retained in KIAA1549::BRAF. **(C)** Representative Airyscan immunofluorescence images of h9-hNSCs expressing C-terminal V5-tagged KIAA1549::BRAF or KIAA1549::BRAF^delTM^ labelled with V5-tag and PDI (marker of endoplasmic reticulum) antibodies. Data are from three independent biological replicates. Quantification was performed by segmenting individual cells, then computing the Pearson correlation coefficient between the V5 and PDI channels for each individual cell. Quantification of V5 and PDI co-localization of V5 and PDI (right). Median Pearson correlation coefficient (K::B = 0.19, K::B^delTM^ = 0.57). Statistical significance was computed with a two-tailed t-test (p < 1×10^−10^). **(D)** Representative images and quantification of colony formation assays of h9-hNSCs expressing Luciferase (control), KIAA1549::BRAF, and KIAA1549::BRAF^delTM^. Data are from three independent biological replicates. Percent area covered: Luciferase = 5.87%, K::B = 89.2%, K::B^delTM^ = 8.78%. Statistical significance was computed with a one-way ANOVA test with Tukey’s multiple comparison test (Luc vs K::B p ≤ 0.0001****, Luc vs K::B^delTM^ p > 0.05). **(E)** Representative immunoblot of pERK, ERK, and Actin in h9-hNSCs expressing the same constructs as in (c) and (d). Quantification of immunoblots in (**Supplemental Fig. 6**). **(F)** Representative images and quantification of colony formation assays of h9-hNSCs expressing Luciferase (control), KIAA1549::BRAF, and KIAA1549::BRAF^ΔSP^. Data are from two independent biological replicates. Percent area covered: Luciferase = 1.2%, K::B = 61.8%, K::B^delSP^ = 0.89%.

The pattern of post-translational modifications to wildtype KIAA1549 supports this model of a single transmembrane domain. Phosphorylation occurs primarily within the cytoplasm, with few exceptions37, whereas glycosylation tends to be extracellular. A single transmembrane domain model would predict that all residues that are C-terminal to the TM domain (residues 1299-1319) would be intracellular and thus more likely to be phosphorylated. Publicly available datasets38 and databases39, show that wildtype KIAA1549 is primarily phosphorylated within residues 1388-1914 of the C-terminal domain (Fig. 4A). Conversely, N-linked glycans have been identified at residues of mouse Kiaa154940 that correspond to 1088, 1095, 1124 and 1226 in human KIAA1549 and residue 1063 exclusively in human KIAA1549 (Fig. 4A). We also previously found that human KIAA1549 is heavily glycosylated (>60 sites) by O-linked mannose (O-Man) glycans within residues 73-86441 (Fig. 4A). Both N- and O-Man glycosylation is initiated within the lumen of the endoplasmic reticulum prior to trafficking of transmembrane proteins to the cell surface. The identification of ER-signal peptide (residues 1-61) and glycosylation within residues 73-1212 of KIAA1549 thus shows extracellular orientation for these residues. Taken together, these data demonstrate that KIAA1549 is a type I transmembrane protein with a predicted trans-membrane domain at residues 1299-1319 (Fig. 4A).

The observation that KIAA1549 harbors a TM domain suggests that oncogenic activity may result from KIAA1549 tethering the BRAF kinase domain to the cytoplasmic face of membranes. We therefore examined the subcellular localization of K::B using immunofluorescence microscopy. Surprisingly, K::B appeared to be distributed throughout the cell, with expression detected in the cytoplasm and at cellular membranes (Fig. 4C). In contrast, K::B in which the putative KIAA1549 trans-membrane domain had been deleted (K::BdelTM) co-localized with the endoplasmic reticulum (ER) marker PDI, indicating that loss of the trans-membrane domain results accumulation in the ER (Fig. 4C, Pearson correlation coefficient K::B = 0.19, K::BdelTM = 0.57, p < 1×10-10). To test whether ER-localized K::B remains sufficient for oncogenic signaling, we assessed two-dimensional colony formation in the absence of exogenous EGF and bFGF (Fig. 4D). While K::B was sufficient to activate ERK (p = 0.0051, K::B vs K::BdelTM) and induce growth under these conditions, K::BdelTM was not (Fig. 4E and Supplemental Fig. 6, p = 0.616).

These data indicate that K::B is trafficked through the ER, in contrast to wildtype BRAF, which is translated directly into the cytoplasm42. Proteins which traffic through the ER often have an N-terminal signal peptide that directs the nascent ribosome-bound peptide to the translocon channel, where it is subsequently translated directly into the ER lumen. To test the necessity of trafficking into the ER, we deleted the KIAA1549 signal peptide (K::BdelSP) and measured the ability of K::BdelSP to transform cells. Again, K::B was sufficient to transform cells whereas K::BdelSP was not (Fig. 4F, 69-fold decrease relative to K::B). The findings that both K::BdelTM and K::BdelSP are non-functional suggest that K::B trafficking through the ER lumen is required for its activity. It also suggests that proteins which facilitate this trafficking may represent dependencies in K::B-driven tumors.

### K::B is subjected to proteolytic cleavage during secretory pathway maturation

We were next interested in understanding additional post-translational modifications beyond the ER that facilitate K::B oncogenic activity. Intriguingly, when we previously had deleted the K::B transmembrane domain, we noted the disappearance of a low-molecular-weight (~110 kDa) band in immunoblots of C-terminally tagged K::B (**Fig. 5A and Supplemental Fig. 7A**). This was accompanied by the accumulation of K::B in the ER on immunofluorescence (**Fig. 2C**). We hypothesized that this band reflects a K::B cleavage product and that trafficking of K::B beyond the ER is necessary for its cleavage. To assess the generalizability of this proposed cleavage event across cell lineages, species, and fusion break-point isoforms, we immunoblotted for the C-terminal HA tag in nine cell lines: SV40 large T antigen-immortalized normal human astrocytes (NHA), HEK-293T cells, and mouse embryonic fibroblasts (MEF) that were each engineered to express K::B^15::9^, K::B^16::9^, or K::B^16::11^. Similar to our finding in K::B-expressing h9-NSCs, a ~110 kDa product was detected across all of these conditions (**Supplemental Fig. 7B**).

**Figure 5.**
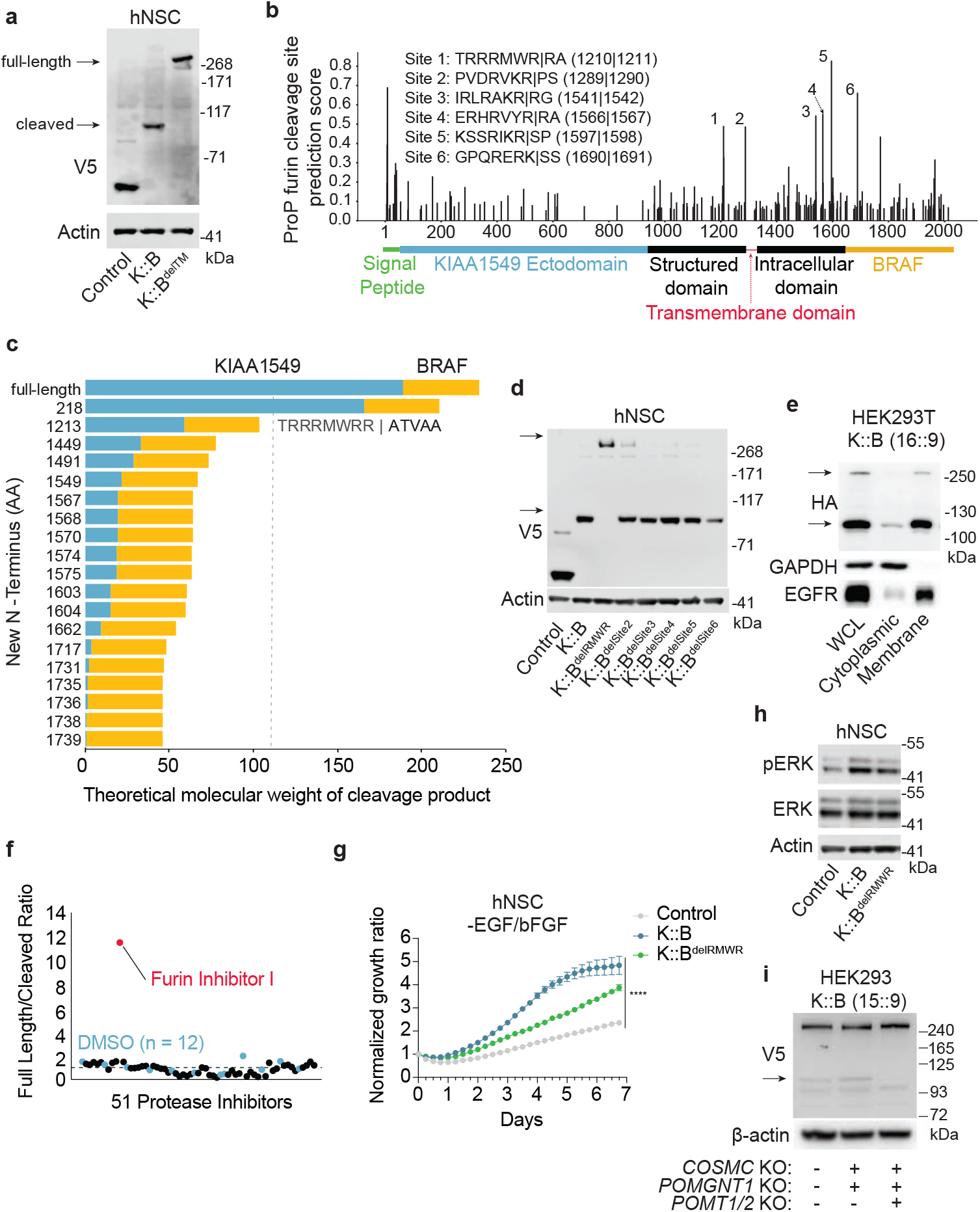
KIAA1549::BRAF is cleaved within the secretory pathway. **(A)** Representative immunoblots showing molecular weight and protein levels of h9-NSCs engineered to express V5-tagged Luciferase (control), KIAA1549::BRAF (K::B), and KIAA1549::BRAF^delTM^ (K::B^delTM^). Arrow depicts a cleaved product within K::B that is not present when the K::B transmembrane domain is deleted (K::B^delTM^). Quantification of three independent replicates is shown in (**Supplemental Fig. 7A**). **(B)** Putative Furin cleavage sites in KIAA1549::BRAF were identified using ProP 1.0 (as described in Methods). The x-axis depicts the location the predicted cleavage site (residue number in KIAA1549::BRAF) and the domain architecture of KIAA1549::BRAF (15::9). The y-axis depicts the ProP prediction score (**Supplemental 30)**. Significant peaks (n = 6) are annotated, and the corresponding context sequence for each cleavage site is displayed in the inset text. **(C)** Bar graph represented the estimated molecular weight of KIAA1549::BRAF (16::9) cleavage products for each peptide identified with Terminal amine isotopic labelling of substrates (TAILS) mass spectrometry. The y-axis depicts the N-terminal residue number for each identified cleavage product. The x-axis depicts the predicted molecular weight of each cleaved product. KIAA1549 residues are displayed in blue and BRAF residues are displayed in yellow. The bar length represents the length of the C-terminal cleavage product for the corresponding identified peptide. The dotted line represents the molecular weight of the 110 kDa band observed in immunoblots of KIAA1549::BRAF-expressing cells (**Fig. 5A**). **(D)** Representative immunoblot (from three independent experiments) showing full-length (top bands) KIAA1549::BRAF and the putative cleavage product (bottom bands) in lysates from h9-hNSCs expressing V5-tagged KIAA1549::BRAF with deletion of predicted Furin cleavage sites 1-6 (See (b) and quantification in (**Supplemental Fig. 7C**). **(E)** Representative immunoblot (from two independent experiments) showing full-length (top bands) KIAA1549::BRAF and proposed cleavage product (bottom bands) in fractionated lysates from HEK293T cells expressing C-terminal HA-tagged KIAA1549::BRAF (16/9). Blots showing whole-cell lysate (WCL), cytoplasmic, and membrane fractions with controls GAPDH (cytoplasmic) and EGFR (membrane). **(F)** A library of 53 protease inhibitors (**Supplemental Table 7**) was screened to identify those which block KIAA1549::BRAF cleavage. The extent of cleavage was assessed with immunoblotting (**Supplemental Fig. 7F)**. Plot depicts quantification of the ratio between full length KIAA1549::BRAF and the putative 110 kDa cleavage product. Values above 1 indicate enrichment of full length KIAA1549::BRAF relative to the cleavage product. A single protease inhibitor (Furin Inhibitor I) was sufficient to block KIAA1549::BRAF cleavage and is highlighted in red. DMSO controls (n = 12) are highlighted in blue **(G)** Incucyte growth curve from one representative replicate of three independent biological replicates. H9-NSCs were transduced with Luciferase, KIAA1549::BRAF, or KIAA1549::BRAF^delRMWR^ and cells were grown for 6.75 days in the *absence* of exogenous EGF and bFGF supplementation. The x-axis depicts the time since the initial image, and the y-axis represents the relative cell confluence (normalized to confluence at 0h). Error bars represent Standard Error of the Mean (SEM). Statistics were calculated with two-way ANOVA across three biological replicates with Tukey’s multiple comparison test (Control vs K::B, Control vs K::B^delRMWR^ and K::B vs K::B^delRMWR^): p ∷ 0.0001**** (**Supplemental Fig 7G-H**). **(H)** Representative immunoblots from three independent biological replicates. Cells were transduced with Luciferase, KIAA1549::BRAF, or KIAA1549::BRAF^delRMWR^ and were grown in the absence of EGF/bFGF for 48 hours before immunoblotting for V5, pERK, ERK, and Actin. Quantification is shown in **(Supplemental Fig. 7I)**. (**I**) Immunoblot depicting V5-tagged KIAA1549::BRAF (top: full length, band 2: cleaved product) expressed in wildtype HEK293, HEK293s with combined knockout of *COSMC/POMGNT1* and a third condition with the addition of *POMT1/2* knockout. Quantification of immunoblot is shown in (**Supplemental Fig. 7J)**

The K::B fusion protein is predicted to harbor six RXXR consensus cleavage sites, which are the preferred substrates for nine PCSK family members including Furin. Five such sites are located within KIAA1549 and a sixth site is in BRAF (**Fig. 5B**). Of the two sites on the luminal side of the trans-membrane domain, one is predicted to be in a flexible loop within a domain that is structurally similar to a SEA domain. The second putative cleavage site is directly adjacent to the transmembrane domain (**Extended Fig 3 and Supplemental Fig. 8A**). SEA domains were first associated with heavily O-glycosylated proteins^59^ and have since been found on a wide variety of proteins, including the mucin MUC1, dystroglycan, and Notch (**Supplemental Fig. 8B**)^36^. A feature of SEA domains is a proteolytic cleavage site between β-strands 2 & 3, either as an autoproteolytic site (with motif GSXXX, where X = hydrophobic, e.g., DAG1 and MUC1/3/12 and MUC17^60^), or as a PCSK/Furin cleavage site (e.g., Notch or GPR126^36,61,62^). Structural homology searches of the AlphaFold-predicted KIAA1549 structure revealed three folded domains, each of which matched both ferredoxin and SEA domains (which are themselves related^36^). None of these three KIAA1549 domains harbor the autoproteolytic cleavage motif. Intriguingly, the most C-terminal of these domains (domain 3) has a PCSK/Furin cleavage motif (^1208^RMWR^1211^, in a loop between α-helix 2 and β-strand 2) (**Supplemental Fig. 8A**). This cleavage motif is conserved in homologous KIAA1549 sequences from human to placozoa, indicating its ancient origins (**Supplemental Fig. 8C**).

We took four approaches to determine the precise site where K::B is cleaved. *First*, we performed Terminal Amine Isotopic Labeling of Substrates (TAILS). This enrichment strategy covalently tags neo-N termini that form from proteolytic cleavage and enables their subsequent isolation and detection with mass spectrometry. This analysis detected 25 peptides: 6 that could be generated by cleavage within BRAF and 19 that could be generated by cleavage within KIAA1549 (**Supplemental Table 6**). Only one of these (ATVAA-GNSVVQVVNVSR) corresponded to the molecular weight of the ~110 kDa putative cleaved product from our immunoblot analysis (**Fig. 5C**). The amino acid sequence upstream of to this cleaved product includes a consensus PCSK/Furin RXXR cleavage site. The precise product places it immediately adjacent to ^1208^RMWR^1211^. *Second*, we transduced H9 NSCs with K::B variants in which each of the six individual cleavage sites were deleted and then measured the ratio of cleaved (~110 kDa) and uncleaved (>250 kDa) protein species by immunoblotting. Deleting the residues corresponding to the ^1208^RMWR^1211^ cleavage site was sufficient to block K::B cleavage (**Fig. 5D, Supplemental Fig. 7C**, K::B vs K::B^delSite1^ p < 0.0001). *Third*, we simultaneously mutated the arginines at positions 1211 and 1212 to alanines in both wildtype KIAA1549 and K::B expressed in HEK293T cells. These residues lie within and adjacent to the ^1208^RMWR^1211^ cleavage motif. This mutation eliminated the ~110 kDa band in KIAA1549 immunoblots (**Supplemental Fig. 7D**). *Fourth*, we performed subcellular fraction experiments of HEK293T cells transduced to express C-terminally tagged KIAA1549 or K::B. Cleavage of KIAA1549 and K::B at ^1208^RMWR^1211^ would be predicted to affect the luminal but not cytoplasmic portion of KIAA1549. Indeed, the 110 kDa cleaved band appeared predominantly in the membrane fractions, consistent with luminal cleavage. (Fig. 5E Supplemental Fig. 7E). We conclude that K::B cleavage occurs within ^1208^RMWR^1211^.

Furin and the other six PCSK isoforms cleave adjacent to an RXXR motif, all of which are inhibited by Furin Inhibitor I^63,64^. To identify candidate proteases that regulate cleavage of K::B, we screened a library of 51 annotated protease inhibitors (**Supplemental Table 7**) and assessing which compounds were sufficient to block K::B cleavage. Only Furin Inhibitor I block K::B cleavage (**Fig. 5F** and **Supplemental Fig. 7F**). Taken together, these data suggest that either Furin or other members of the PCSK family contribute to K::B cleavage at ^1208^RMWR^1211^.

### K::B cleavage requires POMT and enhances MAPK activity

We next hypothesized that proteolytic cleavage of K::B enhances its oncogenic activity, particularly through enhanced MAPK activation by the C-terminal fragment that includes the BRAF kinase. We therefore expressed either full-length K::B or an R^1208^MWR^1211^ deletion mutant (K::B^delRMWR^) in h9-NSCs and measured growth kinetics in the presence or absence of exogenous EGF and bFGF supplementation (**Fig. 5G and Supplemental Fig. 7G-H**). Expression of either construct induced ERK phosphorylation and growth-factor independent growth (**Fig. 5G-H**), but less so in the deletion (p = 0.017 and p < 0.0001, respectively) (**Fig. 5G-H, Supplemental Fig. 7I**). We conclude that cleavage at R^1208^MWR^1211^ enhances K::B oncogenic activity through the MAPK pathway.

K::B cleavage requires *POMT1/2* expression. We expressed K::B in both wild-type HEK293 cells, HEK293 cells with deletion of the O-glycosylation enzymes *COSMC* and *POMGNT1*, and HEK293 cells with deletion of *POMT1/2* (double knockout) in addition to *COSMC* and *POMGNT1*. We observed the ~110 kDa C-terminal K::B fragment in both wild-type and the *COSMC/POMGNT1* deletion mutants, but not in the context of *POMT1/2* deletion (p = 0.04) (**Fig. 5I and Supplemental Fig 7J**).

We conclude that K::B fusions are associated with a cleavage product across multiple contexts and that proteolytic maturation depends on POMT1/2 mediated O-mannosylation in the ER.

### Pomt1 is required for KIAA1549::BRAF fusion activity *in vivo*

The finding of POMT1/2 dependency across our *in vitro* models raises the possibility that it may represent a potential therapeutic target for children with K::B-driven pLGGs. We thus evaluated whether *Pomt1* suppression reverses *in vivo* phenotypes associated with K::B expression in the murine brain. To generate *in vivo* models of K::B expression, we leveraged in-utero-electroporation (IUE) of embryonic mouse brains. IUE of the 16::9 variant of the K::B fusion into the embryonic day 14.5 (E14.5) cortex resulted in abnormal migration of GFP-positive transfected glutamatergic neurons and increased glial reactivity seen by Gfap expression in both transfected and non-transfected cells compared to mice subjected to IUE of vector control (Fig. 6A, Supplemental Fig. 9A). The increased glial reactivity in K::B-expressing cells resulted in striking differences in astrocytic morphology. While vector control transfected cortical gray matter astrocytes displayed classic, finely branched protoplasmic morphologies, those in the K::B condition showed dense and ramified morphologies associated with fibrous and reactive astrocytes (**Fig. 6A, Supplemental Fig. 9A**). These observations suggest that expression of K::B in embryonic neural stem cells induces developmental and morphological changes in subsequently produced neurons and glia that persists into adulthood.

**Figure 6.**
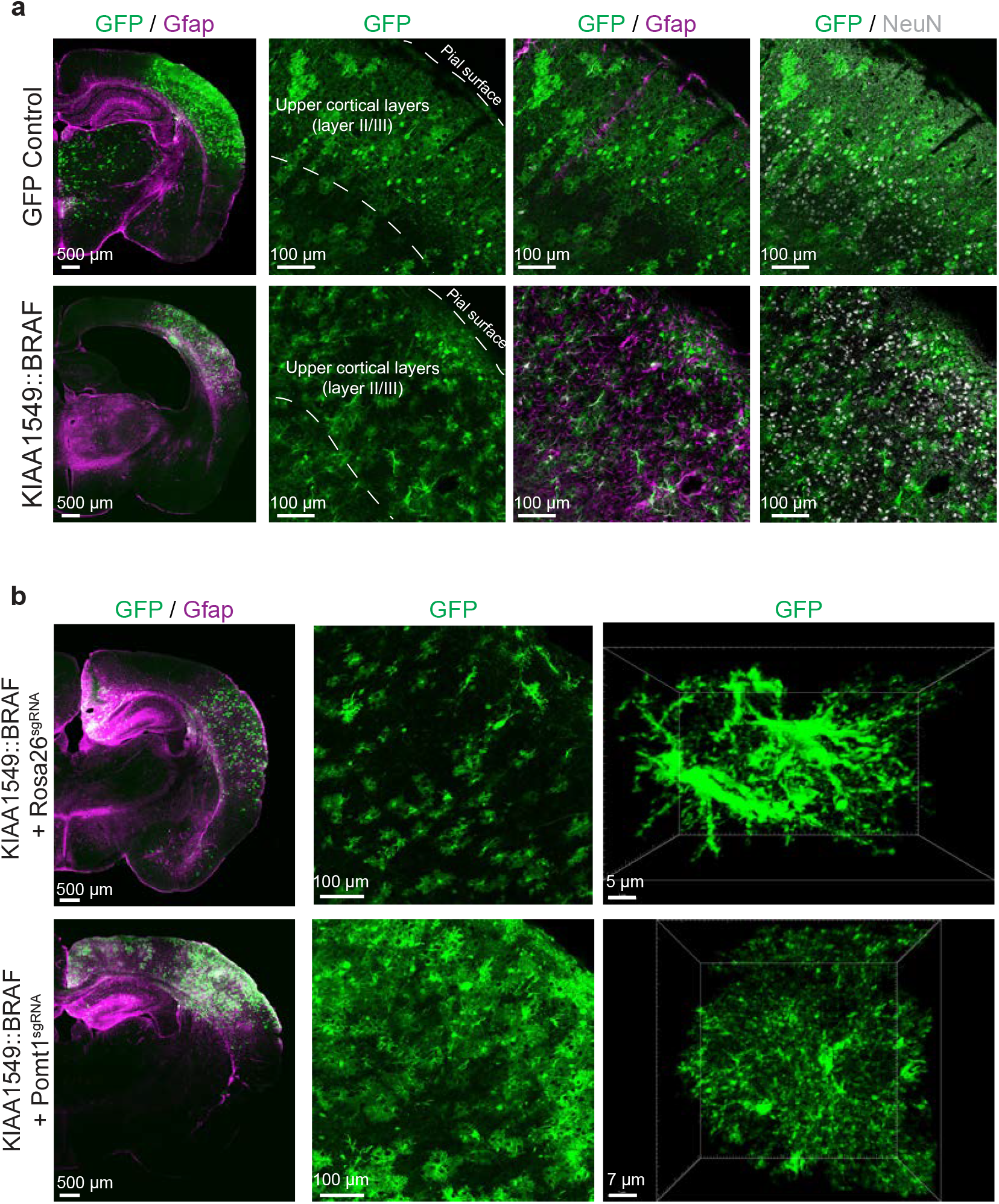
Pomt1 suppression reverses in vivo brain phenotype changes induced by KIAA1549::BRAF expression. **(A)** Representative images depicting EGFP+ transfected cells within EGFP control and KIAA1549::BRAF expressing brain sections (left). High magnification representative images of the upper cortical layer region co-stained for EGFP (middle left, green), EGFP/Gfap (middle right, green and magenta) or NeuN (right, green and white/grey). **(B)** Representative images depicting EGFP+ transfected cells within K::B + Rosa26 sgRNA and K::B + Pomt1 sgRNA expressing brain sections. High magnification representative images of the upper cortical layer region depicting EGFP+ cell morphology (middle) and 3D reconstruction of example EGFP+ astrocytes in each condition (right). Scale bars: tiled section images on left side of (A) and (B) = 500 μm; high magnification images = 100 μm; 3D reconstruction images = 5 μm (K::B + Rosa26 sgRNA) and 7 μm K::B + Pomt1 sgRNA.

With this model, we next evaluated whether *Pomt1* suppression reverses K::B-associated *in vivo* phenotypes. We first co-electroporated CRISPR/Cas9 components targeting *Rosa26* (control *Rosa26*^sgRNA^) or *Pomt1* (*Pomt1*^sgRNA^) with plasmids expressing the K::B fusion and EGFP for visualization and validated efficiency of *Rosa26*^sgRNA^ and *Pomt1*^sgRNA^ by isolating and culturing neural stem cells 24 hours post-IUE with EGFP and sgRNA/Cas9 components. *Rosa26*^sgRNA^- and *Pomt1*^sgRNA-^electroporated cells showed indel rates above 70% and 90%, respectively (**Supplemental Fig. 9C**). K::B + Rosa26^sgRNA^-electroporated brains displayed phenotypes similar to prior K::B fusion samples, including mislocalization of transfected neurons and glial reactivity within the electroporated region, with gray matter cortical astrocytes displaying reactive and fibrous morphology (**Fig. 6B**). These reactive astrocytes displayed little to moderate branching and had clearly distinguishable thickened truncal processes. However, brains harvested from K::B + *Pomt1*^sgRNA^ contained transfected cells that more closely resembled those found in EGFP control-electroporated brains. Specifically, most EGFP-positive transfected K::B + *Pomt1*^sgRNA^ astrocytes displayed classic protoplasmic, spongy astrocyte morphology, with many fine branches and branchlets, similar to EGFP-control brains and in stark contrast to those found in K::B + Rosa26^sgRNA^ brains (**Fig. 6B**). Taken together, we conclude that expression of K::B induces developmental and morphological changes in the brain that require *Pomt1/2* for their manifestation.

## Discussion

These findings provide the first proof of concept that fusions can be therapeutically targeted beyond the proto-oncogene. Contrary to current dogma, we find that BRAF fusion partners are necessary for transformation and induce novel vulnerabilities. In the setting of K::B, the KIAA1549 fusion partner engenders striking and specific dependency on the POMT complex, which is necessary for its protein maturation to facilitate oncogenesis.

We show that K::B fusions undergo a series of post-translational modifications that contribute to the intracellular trafficking and activation of the K::B proto-oncogene that are required for oncogenesis, thereby engendering novel vulnerabilities. The results of our experiments with KIAA1549 and FAM131B indicate that these BRAF fusion partners are essential for the oncogenic function of their BRAF fusions and suggest that this property extends to other BRAF fusion partners. It is likely that other fusion oncogenes also undergo post-translational modifications and intracellular trafficking that promote oncogenesis, and that these may afford novel dependencies in other cancer types. Other oncogenic fusions, including FGFR fusions, likely also traffic through the secretory pathway given they retain the FGFR transmembrane domain; the role of glycosylation in these contexts is similarly unclear.

Our findings stand in contrast to a prevalent view that truncated BRAF kinase alone is sufficient for transformation, based on experiments that have focused on the BRAF kinase^8,65–68^. However, all prior studies that showed *in vitro* transformation phenotypes due to expressing the truncated BRAF kinase included additional N-terminal amino acids that are not in the wild-type protein, or were performed in immortalized cells^65,67,69,70^. The only prior *in vivo* report in wildtype mice found that truncated BRAF alone is insufficient to induce pilocytic astrocytoma formation^71^, consistent with our findings.

It is possible that anti-transformation screens of the sort we performed with K::B might help identify co-factors that enable necessary post-translational modifications and trafficking of other fusion oncoproteins. Extensive efforts have been directed towards leveraging synthetic lethal screens to detect potential therapeutic targets whose modification is synthetic lethal with common genetic alterations in cancer. These include prominent successes such as PARP1 inhibition in the context of BRCA1/2 loss^72^, PRMT5 suppression in the context of MTAP deletions^73–75^, and WRN suppression in the context of microsatellite instability^76,77^. Our screening approach was capable of identifying genes whose loss is synthetic lethal with K::B expression, but also assessed for reversal of a transformation phenotype (growth in the absence of EGF/bFGF). The design of this screen also lends itself to the use of isogenic controls, which can limit the effects of confounding variables.

The function of KIAA1549 remains unknown, but its properties of heavy mannosylation and cleavage suggest similarities to signaling mucins. Signaling mucins are a diverse class of proteins that share many common architectural characteristics (densely O-glycosylated extracellular domain and a cytoplasmic signaling domain), but differ in the number and nature of key details such as the type of glycosylation found on the extracellular domain, the presence of SEA domains, and the presence/effect of extracellular cleavage^78,79^. An example of a prototypical animal signaling mucin is MUC1, whose extracellular mucin domain carries O-GalNAc-type glycosylation, possesses an autoproteolytically cleaved SEA domain, and has a cytoplasmic tail for signaling^80^. In contrast to this mucin, baker’s yeast (Saccharomyces cerevisiae), encodes for two *bona fide* O-Man signaling mucins – Msb2 and Hkr1, both of which regulate MAPK signaling^81^. Moreover, and similarly to K::B, knockout of the O-mannosyltransferase Pmt4 regulates MAPK signaling^82^. Both Pmt4 and POMT1 belong to the PMT4 subfamily, sharing similar capacity for O-glycosylation of substrates^83^. Furthermore, Msb2 possesses a cleavage domain between residues 1045 and 1145, whose cleavage by Yps1 is required for MAPK activation^84^, and overlaps with an unannotated SEA domain.

KIAA1549 and K::B fusions share features with the classical POMT1/2 substrate DAG1, whose functions also depend on O-mannosylation for functions^85^. Autoproteolysis in the SEA domain of DAG1 facilitates maturation into αDG and βDG chains. Mutants that disrupt DAG1 SEA domain cleavage traffic to the cell surface, acquire functional O-mannosylation, and retain laminin binding properties^86^ — indicating that SEA domain cleavage is dispensable for DAG1 functions. However, we do not yet know whether wild-type KIAA1549 or K::B depend upon SEA domain cleavage for function.

Our findings harbor particular significance for therapeutic approaches for children with K::B driven pLGGs and the occasional extracranial tumors with K::B fusions^9,87^. The last decade has seen unprecedented progress with the use of MAPK pathway inhibitors as a therapeutic approach for these children, culminating in the recent FDA approval of the pan-RAF inhibitor tovorafenib^14^. However, approximately 30% of K::B driven gliomas are innately resistant. Those that respond often exhibit rebound growth upon cessation of therapy, necessitating continuous long-term treatment. Long-term treatment can be intolerable: the MAPK pathway is essential for the development of many tissues and organs, and children have experienced severe reductions in growth velocity while on treatment^13,88^.

These considerations increase the value of alternative strategies to target K::B fusions, such as inhibiting POMT1/2, the DPM complex, or PCSK proteases. Indeed, whereas MAPK inhibitors may only induce cell cycle arrest or senescence^89^ targeting K::B trafficking may have the additional benefit of causing toxic accumulation of K::B within the secretory pathway. This would recall the sensitivity of transformed B-cells (which produce high levels of immunoglobulin) to proteosome inhibitors, which lead to the toxic accumulation of immunoglobulin in the ER^90^. Targeting POMT1/2 may result in its own toxicities: congenital homozygous loss-of-function mutations have been associated with several syndromes, most prominently muscular dystrophies^91–96^. This likely reflects effects on DAG1^41,47,48,93^. However, heterozygous carriers of these mutations exhibit few symptoms^93,96^, raising the possibility that partial inhibition of the POMT complex in postnatal tissue may be tolerated—perhaps in combination with MAPK pathway inhibition.

Finally, our observation that the N terminus of K::B is heavily mannosylated and likely secreted raises an intriguing possibility that this is the mechanism causing the formation of tumor-associated mucinous cysts, which are pathognomonic to PAs and currently only treatable through surgery^10^. Mucinous proteins have a preponderance of O-GalNAc adducts; while different from O-mannosylation, it raises the possibility that heavily glycosylated KIAA1549 ectodomain may be secreted, contributing to cyst formation. Disrupting K::B trafficking and thus cleavage would thereby be expected to reduce KIAA1549 ectodomain shedding and cyst formation. The identification of the O-mannosylation complexes as potential MAPK-independent therapeutic targets is therefore of substantial clinical importance.

## Acknowledgements

We would like to express our sincere thanks and gratitude to the children and their families who have made this study possible by participating in tumor banking protocols and advocating for and supporting research for pediatric low-grade gliomas. We would also like to thank the following organizations and foundations for their invaluable support and contributions to our research: The Dana-Farber Pediatric Low Grade Glioma Program: PB, RB, KLL, EF, ME, SM, AB. PLGA Fund at the Pediatric Brain Tumor Foundation: PB, RB, KLL, SM, AB. Team Jack Foundation: PB, RB, TP, DTWJ. The Jared Branfman Sunflowers for Life Fund: PB, RB. Helen Gurley Brown Foundation: PB, AB. Lauren’s First and Goal Foundation: PB, RB. The Everest Centre for Low-grade Paediatric Brain Tumours: DTWJ, TM. Jack’s Drive 55: PB, RB. Friends of DFCI: PB, RB. The Brain Tumour Charity: PB, RB. The Gray Matters Brain Tumor Foundation: RB. Broad Institute Escape Velocity Award: PB. Alex’s Lemonade Stand Foundation: SM. Swedish Childhood Cancer Foundation: AB. Swedish Research Council: AB. Pedals for Pediatrics: PB, AB. David M. Livingston, MD, Physician-Scientist of the Damon Runyon Cancer Research Foundation, and an Edward P. Evans Foundation Evans MDS Young Investigator: MV. Novo Nordisk Foundation (NNF22OC0076899): HJJ. The Carlsberg Foundation and a research grant (00025438) from VILLUM FONDEN: AH. German Cancer Consortium (DKTK) joint funding pool project Next Gen LOGGIC: TM and TB. D-A-CH funding scheme between the German Research Foundation and the Swiss National Science Foundation SNF: JD and TB. DFG funded Heisenberg professorship: TB. Canton and University of Fribourg as a member of the SKINTEGRITY.CH research network: JD.

We would like to thank Kun Huang from the DFCI Molecular Imaging Core for assisting with this study, and Drs. Charles Stiles and David Misek for helpful discussion.

## Conflict of interest

PB has received grant funding from Novartis Institute of Biomedical Research, and has served on paid advisory boards for QED Therapeutics, and Day One Biopharmaceuticals. RB consults for and owns equity in Scorpion Therapeutics, Karyoverse Therapeutics, and LOH Therapeutics. WCH is a consultant for Thermo Fischer, Solasta Ventures, KSQ Therapeutics, Frontier Medicines, Jubilant Therapeutics, RAPPTA Therapeutics, Serinus Biosciences, Kestral Therapeutics, Crane Biotherapeutics, Function Oncology, Recursion Pharma, and Weaver Biotherapetics and Perceptive. DER receives research funding from members of the Functional Genomics Consortium (Abbvie, BMS, Jannsen, Merck, Vir), and is a director of Addgene, Inc. TM is advisory board member for Ipsen Pharma GmbH. SAM is a consultant for Karyoverse Therapeutics and is a consultant for and owns equity in LOH Therapeutics. ESF is a founder, scientific advisory board (SAB) member, and equity holder of Civetta Therapeutics, Proximity Therapeutics, Neomorph, Inc. (also board of directors), StelexisBiosciences, Inc., Anvia Therapeutics, Inc. (also board of directors) and CPD4, Inc. (also board of directors). He is an equity holder and SAB member for Avilar Therapeutics, PhotysTherapeutics, and Ajax Therapeutics and an equity holder in Lighthorse Therapeutics. E.S.F. is a consultant to Novartis, EcoR1 capital, Odyssey and Deerfield. The Fischer lab receives or has received research funding from Deerfield, Novartis, Ajax, Interline, Bayer and Astellas. TB received speaker honoraria from Pierre Fabre and the European Society for Medical Oncology (ESMO)

## Author contributions

SM, AB, DC, GK, RB, HJJ, TB, AH and PB conceived of the study. AZ, SR, MB, KC, KZ, EMG, AG, LLV, TJR, ZK, NP, TA, EM, SV, MS, JJ, SMA, LE, JP, ALC, AF, SHC, MO, ZE, JL, DN, SB, AA, JR, KT, AM, DB, JK, MV, HJ, SP, TG, WCH, EF, JSB, JD, HC, NJ, TM, AC, BAC, KL, KJ, ME, DR, DTWJ and TNP contributed by assisting with performing and/or supervising the experiments. SM, AB, DC, GK, RB, HJJ, TB, AH and PB wrote the manuscript. All authors reviewed, edited, and approved the manuscript. RB, HJJ, TB, AH and PB were responsible for overseeing the overall project.

## Additional information

Supplementary Information is available for this paper. Correspondence and requests for materials should be addressed to the corresponding authors.

## Code Availability

All code for this manuscript is deposited in the following github repository: https://github.com/beroukhim-lab/pomt_manuscript_code.

## Data Availability

The mass spectrometry glycoproteomics data have been deposited to the ProteomeXchange Consortium via the PRIDE^97^ partner repository with the dataset identifier PXD062030. The mass spectrometry TAILS proteomics data have been deposited to the ProteomeXchange Consortium via the PRIDE^97^ partner repository with the dataset identifier PXD062076. DepMap data (24q2) were downloaded from the depmap.org web portal.

## Methods

### Patient sample collection

Patient-derived pediatric low-grade glioma samples were obtained from the Dana-Farber/Boston Children’s Cancer Center Brain Tumor Bank, governed by the Dana-Farber IRB approved protocol DFCI 10417. Informed consent for tumor banking was provided by patients and/or their guardians. Additional samples were obtained McGill University and collected under an institute approved protocol with full informed consent.

### Analysis of DFCI sequencing cohort

An initial group of adult and pediatric neuro-oncology tumors was identified from a DFCI institutional database (OncDRS) followed by a query from two CNS-tumor specific REDCap databases that have undergone IRB approval (10-043 and 10-417 REDCaps). A cohort of >5000 tumors was identified, all of which had targeted exome sequencing via OncoPanel performed by the Brigham and Women’s Hospital Center for Advanced Molecular Diagnostics (BWH CAMD) for at least one of their brain tumor samples (N = 5115 tumors). Patients were deemed eligible if their first pathologic brain tumor diagnosis was at the age of 21 years or younger and they had a pathologically confirmed pediatric low grade glioma diagnosis (N = 435 tumors). Due to the diversity of pediatric low grade glioma diagnoses (pLGG) and the increasing reliance on molecular characterization to finalize a pLGG diagnosis, tumor diagnoses were given in a stepwise manner. Following the WHO 2021 Classification of Central Nervous System Tumors, tumors with a corresponding top line diagnosis and molecular characteristic (e.g. pilocytic astrocytoma with BRAF rearrangement or ganglioglioma with BRAF^V600E^ mutation) were classified as such. Similarly, tumors with a top line diagnosis of low-grade glioma, not otherwise specified (LGG, NOS) with a single MAPK alteration were manually reviewed for histologic features of a corresponding pLGG diagnosis. These tumors were classified more specifically if the necessary histologic features were mentioned in the microscopic description of the pathology report. Any tumors that did not meet the criteria for more specific classification were grouped into the LGG, NOS diagnosis. All deidentified clinical and molecular information was coded in an Excel sheet and converted to a text file for the finalized OncoPrint.

### Single-cell RNA-sequencing analysis

#### scRNAseq tissue collection and processing

Tumor samples obtained from DFCI and McGill University Health Centre. Glioma biopsies were dissociated using Miltenyi, #130-095-942. Cells were loaded Chromium Controller to generate single-cell gel beads in emulsion, before RT-PCR amplification followed by indexed library preparation. An Agilent 2100 Bioanalyzer was used to check the quality control of cDNA library concentration and fragment size before proceeding to NextSeq sequencing, aiming for a minimum depth of ~20,000 reads per cell. During preprocessing, we removed cells in which fewer than 200 genes were detected and filtered out genes with poor coverage (ie they were detected in fewer than three cells). Finally, we applied a threshold of 30% for mitochondrial genes to exclude low-quality cells, and excluded genes including **MALAT1**, mitochondrial genes (**MT-**), and hemoglobin genes (**HB** excluding HBP). Reads were normalized and log transformed. We performed **Principal component analysis (PCA)** using the **arpack** solver, retaining the top **30 components**. Harmony integration was used to correct for batch effects, following which we applied **UMAP** dimensional reduction to visualize cellular heterogeneity.

### Bulk RNA-sequencing analysis

Bulk RNA sequencing data of primary human low-grade glioma samples was generated as previously described^7,98^, with transcript count matrices used for downstream analyses, prior to evaluating expression of *POMT1, POMT2, DPM1, DPM2* and *DPM3* across the dataset.

### Analysis of *POMT1/2* and *DPM1-3* expression in normal adult tissues

Bulk tissue gene expression data (RNA-sequencing) for *POMT1* and *2*, and *DPM1-3* was obtained from the GTEx Portal^50^ on 12/13/2024. The data covers all available tissues from both sexes and is presented as transcripts per million (TPM). The Genotype-Tissue Expression (GTEx) Project was supported by the Common Fund of the Office of the Director of the National Institutes of Health, and by NCI, NHGRI, NHLBI, NIDA, NIMH, and NINDS.

### Fusion partner analysis in pediatric and adult tumors

Pediatric kinase fusion partners were collected as previously described^87^. Inframe fusions from adult tumors were downloaded from FusionGDB 2.0^15^. Fusions were filtered to include only in-frame fusions with annotated breakpoints. The number of unique fusion species, including fusions with different partner genes and those with different intronic breakpoints for the same partner gene, was computed for each fusion partner gene. To map BRAF breakpoint loci, fusions in the pediatric kinase dataset were filtered to include only those in which BRAF is a fusion partner.

### Protein topology, hydrophobicity, myristoylation, and cleavage site analysis

Protein topology prediction of signal peptides, intracellular domains, extra-cellular domains, and transmembrane domains was done with DeepTMHMM^32^. Proteins with a MGXXX(S/T) sequence at positions 1-5 in their coding sequence were annotated as having a consensus myristoylation site as described in. Local hydrophobicity was computed as described in^33^ by binning the KIAA1549 amino acid sequence into 20-mers, then computing the hydrophobicity score using previously described weights^33^. Furin and proprotein convertase cleavage sites were predicted using ProP 1.0^99^.

### Alphafold prediction of KIAA1549 structure

The predicted structure for human KIAA1549 (accession AF-Q9HCM3-F1-v4) was retrieved from the AlphaFold database^35,100^, and paucimannose Nglycans were attached to the structure as illustrative of N-glycosylation using the Re-Glyco tool^101^.

### Genome-scale anti-transformation screens

Genome-scale CRISPR/Cas9 screens were performed in mNSCs transduced to express K::B, BRAF^V600E^ or luciferase controls as previously described^44^. The KIAA1549-BRAF and BRAF^V600E^ screens were performed in the absence of exogenous EGF/bFGF supplementation, thereby rendering cells oncogene dependent and allowing the identification of genes/pathways that are necessary for transformation. Screen analysis was performed by first computing the differential in fold change in CRISPR guide abundance between the early (day 3) and final (day 18) timepoint samples. Single-gene scores were computed by first averaging all four sgRNAs for a given gene, then averaging across biological replicates. To identify statistically significant differences in guide depletion, guide-level depletion scores were collapsed into mean gene-level depletion scores across all four guides, then statistically significant differences across replicates were computed with two-tailed t-tests with Benjamini-Hochberg^102^ multiple hypothesis correction. Depletion scores were compared between the luciferase (control) screen and either the BRAF^V600E^ or KIAA1549::BRAF screens.

### Analysis of gene essentiality across cancer cell lines

Gene essentiality scores for 1150 cell lines profiled in The Dependency Map^54^ 24q2 release are described in **Supplemental 19**. The chronos^55^ gene essentiality scores were plotted for each of six genes, *POMT1, POMT2, DPM1, DPM2, DPM3*, and *POLR2B*, for all 1150 cell lines.

### Antibodies, chemical compounds, primers, CRISPR guides, and plasmid constructs

Antibodies are described in (**Supplemental Table 8**). Primers are described in (**Supplemental Table 9**). CRISPR guides were designed using CRISPick^103–106^ and are described in (**Supplemental Table 10**). Plasmid constructs were synthesized by Epoch Life Science (Houston, USA) or Genscript (Piscataway, USA) and are described in (**Supplemental 38**). R3A-5a (2-[(5Z)-4-oxo-5-[[3-(1-phenylethoxy)-4-(2-phenylethoxy)phenyl]-methylene]-2-thioxo-thiazolidin-3-yl]acetic acid)^57^ was synthesized by Genesis Drug Discovery & Development (Montreal, Canada) and quality control is described in (**Supplemental 39**).

### Tissue culture

#### Culture of h9-NSCs

h9-NSCs (Invitrogen, N7800-100) were cultured on geltrex-coated (ThermoFisher, #A1413302) cell culture plates in growth medium containing 500 mL NeuroBasal A (Gibco, 1#0888022), 500 mL DMEM/F-12 (Gibco, #11330057), 20 mL B-27 (Gibco, #17504044), 10 mL HEPES (ThermoFisher, #15630080), 10 mL Glutamax (ThermoFisher, #35050061), 10 mL Sodium Pyruvate (ThermoFisher, #11360070), 10 mL MEM NEAA (ThermoFisher, #11140050), 10 mL Pen-Strep (ThermoFisher, #15140122), 20 ng/mL EGF (STEMCELL Technologies, #78006.2), 20 ng/mL bFGF (STEMCELL Technologies, #78003.2), and 1 mL of Heparin (STEMCELL Technologies, #07980). This growth medium is herein referred to as TSM. For growth factor withdrawal experiments, cells were cultured in TSM without addition of EGF, bFGF, and Heparin. Cells were detached with accutase, pelleted, and resuspended in TSM to remove residual accutase which disrupts re-attachment of cells when subculturing. Cells were subcultured at a ratio of 1:5 – 1:2 into freshly coated cell culture plates and split when cells reached a confluence of 75-90%. To generate growth factor-independent models, cells were transduced with pDEG-KIAA1549::BRAF or pDEG-BRAF^V600E^ and were cultured in TSM without EGF/bFGF for at least three weeks.

#### Culture of mNSCs

Mouse neural stem cells (mNSCs) were established from the subventricular zone of E14.5 mouse embryos (CD-1) and cultured as neurospheres in TSM in ultra-low attachment treated plates and flasks as previously described^107^. Neurospheres were dissociated with accutase at 37°C for 5-10 minutes when passaging or seeding for experiments. Mouse NSCs were cultured in TSM media. For growth factor withdrawal experiments, cells were cultured in TSM without addition of EGF, bFGF, and Heparin. Neurospheres were dissociated with accutase, pelleted, and resuspended in TSM to remove residual accutase. Cells were seeded at a density of 120,000 cells/ml into ultra-low attachment cell culture flasks and split two times per week. To generate growth factor-independent models, cells were transduced with KIAA1549::BRAF or BRAF^V600E^ and were cultured in TSM without EGF/bFGF for at least three weeks.

#### Culture of HEK-293T cells

HEK-293T cells were grown in DMEM + 10% FBS + 1% Pen/Strep. Cells were subcultured at a 1:10 ratio and were split when cells reached a confluence of 75-90%.

#### Culture of NHA

The SV40 large T antigen (TAg) mediated immortalisation of normal human astrocytes (NHA) and their transduction has been described previously^108,109^. NHAs were grown in astrocyte medium (Lonza).

#### Culture of MEFs

The generation and retroviral transduction of TAg immortalised murine embryonic fibroblasts (MEFs) has been described previously^110^. MEFs were cultivated in DMEM medium (4.5 g/l glucose) supplemented with 10% fetal calf serum, 2 mM L-glutamine, 10 mM HEPES, 200U/ml penicillin, 200 µg/ml streptomycin. Tissue culture media and additives were purchased from PAN Biotech (Aidenbach, Germany).

All cell lines were routinely screened for mycoplasma contamination with a commercially available kit (Lonza, #LT07-318)

### Quantitative PCR (qPCR)

Quantitative PCR was used to validate expression of KIAA1549::BRAF or BRAF^V600E^ in isogenic mouse neural stem cell lines. RNA was isolated using either Qiagen RNeasy Mini kit (Qiagen, #74106) or Qiagen RNeasy Micro kit (Qiagen, #74004). cDNA was made using iScript™ Reverse Transcription Supermix for RT-qPCR (Bio-Rad, #1708840), then used to quantify gene expression with SsoAdvanced Universal SYBR Green Supermix (Bio-Rad, #1725272) on a Quantstudio flex7. All gene expression values were normalized to B2M expression and fold changes were analyzed using the ΔΔCt method. Primers for human *KIAA1549::BRAF* and *BRAF*^V600E^ are described in (**Supplemental Table 9**).

### Lentivirus production and transduction

#### Viral production

HEK293T cells cultured in Dulbecco’s Modified Eagle Medium (Gibco, 11965118) with 10% fetal bovine serum and with or without 1% Penicillin-Streptomycin (ThermoFisher, 15070063). HEK-293T cells were seeded into 10-cm tissue culture plates at a density of 3-5.38×10^6^ cells/plate. The following day cells were transduced with 1 mg of either VSV-G (Addgene Plasmid #14888) or pMD2.G (Addgene Plasmid #12259), 10 mg of psPAX (Addgene Plasmid #12260), and 10 mg of lentiviral plasmid using Lipofectamine 3000 (#L3000015, ThermoFisher, Waltham, USA) according to the manufacturers protocol. The following day the growth medium was changed to 10 mL of TSM^44^ growth medium. Approximately 24 hours later, the growth medium was collected, filtered through a 0.45-micron syringe filter. Viral supernatant was concentrated using LentiX Concentrator (Takara, PT4421-2) according to the manufacturer’s protocol. Alternatively, lentiviral supernatant was concentrated to a final volume of 200 mL with an Amicon 10 kDa centrifugal filter (#UFC9010, SigmaAldrich, St. Louis, USA). Virus was stored at −80 and thawed on ice prior to use.

#### Viral transduction

h9-NSC cells were seeded into geltrex-coated 10-cm plates at a density of 2-3×10^6^ cells/plate. Plates were supplemented with 5-20 mL of concentrated virus per 1×10^6^ cells. The following day the growth medium was changed to 10 mL of TSM^44^ growth medium and, if applicable, selection antibiotics (Puromycin 2 ug/mL or Blasticidin 10-20 ug/mL) were added. Cells were cultured for 5-7 days prior to being used in experiments. Mouse neural stem cells were infected with lentiviral and incubated with centrifugation at 700g at 30°C for two hours. Cells were selected with either 0.5 ug/mL puromycin (for three days) or 10 ug/mL Blasticidin (for 10 days).

### CellTiterGlo Viability assays

h9-NSCs were seeded into geltrex-coated 96-well plates (Corning, 3903) at a density of 1,000-5,000 cells/well in a final volume of 100 mL of TSM growth medium. The next day cells were treated with the indicated concentrations of pomalidomide diluted in an additional 100 mL of TSM growth medium. Cells were grown for the indicated time, then 50 mL of CellTiterGlo (Promega, #G7572) was added to each well. Plates were incubated for 10 minutes at room temperature, then luminescence was measured on an Envision plate reader or a SpectraMax M5 plate reader. Luminescence signal in compound-treated wells was normalized to vehicle control.

Mouse NSCs were seeded into 96-well plates (Thermo Scientific, 165306) at a density of 2,000 cells/well in 90 µL of TSM growth medium. Cells were left for at least 30 mins before treated with 10 µl of medium containing R3A-5a at the indicated concentration (final concentrations: 50, 25, 15, 12, 10, 7.5, 3.75, 0.9375 or 0.234 µM) or equivalent DMSO controls. Cells were grown for 5 days then 100 µL of CellTiterGlo (Promega, #G7572) was added to each well. Plates where incubated on a shaker at room temperature for 10 mins, then luminescence was measured on an Envision plate reader or a SpectraMax M5 plate reader. Luminescence signal in compound-treated wells was normalized to vehicle control. Area under the curve (AUC) was calculated for each experimental replicate (n = 3) and compared using oneway ANOVA with Tukey’s multiple comparison test in Graphpad Prism Version 10.4.1.

### Colony formation assays

Cells were plated in geltrex-coated 6-well plates at a density of 5-10×105 cells/well in a final volume of 2 mL TSM without growth factors. Growth medium was changed every 2-3 days. After 21 days cells were fixed with 3.7% paraformaldehyde for 15 minutes at room temperature and then stained with a 1% Crystal Violet solution (w/v) for 1 hour at room temperature. The 6 well plates were rinsed in ultrapure water, then dried for 24 hours prior to imaging on either a flatbed scanner or the LiCOR Odyssey M.

### Incucyte growth experiments

Human h9-hNSCs were seeded at a density of 20,000 cells/well in geltrex coated 48-well plate (Corning, #3548) in TSM growth media with or without EGF/bFGF supplementation and with or without pomalidomide (1 ug/ml). Plates were placed in the IncuCyte S3, set for imaging every six hours, four images/well with 10x objective. We then analyzed confluence with IncuCyte Basic Analyzer, AI Confluence with segmentation set to default and normalized to the 0-hour timepoint.

Mouse NSCs were seeded at a density of 1,000 cells/well in u-bottom ultra-low attachment 96-well plates (Corning, #3548) in TSM growth media with or without EGF/bFGF supplementation. Cell plates were centrifuged at 200g for 10 mins at room temperature and then placed in the IncuCyte S3, set for imaging every six hours, 4x objective. We then analyzed largest brightfield object area with IncuCyte Spheroid, with segmentation: Sensitivity: 10, Area ≥ 2,000 µm^2^, Eccentricity ≤ 0.75.

### Pomt1 and Pomt2 knockout and cumulative doublings assays

Mouse NSCs were plated as 2×106 cells/well in 2 mL TSM growth media with (Luciferase control) or without (KIAA1549::BRAF) EGF/bFGF supplementation in 12-well plates. Concentrated lentiCRISPRv2 lentivirus (15 µL) encoding both Cas9 and sgRNA targeting mouse Pomt1/2, two guides per target (sgRNA sequences in) as well as EGFP was added to each well in four technical reps. The plates where centrifuged for two hours, 700g at 30°C and then left overnight in incubator. The day after the cells were centrifuged at 340g for 4 mins, virus media was aspirated, and cells were replated in 20 mL fresh TSM growth media with/without EGF/bFGF. 1 ug/mL Puromycin was used for selection over 72 hours after which cells were counted on a Countess 3 FL Automated Cell Counter (Thermo Fisher Scientific). Cells were replated in fresh growth media (500,000 in 5 ml media) in T25 ultra-low attachment flasks. After this, cells were passaged and counted every 3-4 days until the 18-day post transduction endpoint. Each time the same reseeding conditions were used. Cumulative doublings were calculated using end of selection as starting point and with the formula log2(total viable cells/seeded viable cells).

### CRISPR/Cas9 competition assays

Cas9 guide sequences used in CRISPR competition assays are described in (Supplemental Table 10). Negative control (intergenic) guides were cloned into an EGFP-expressing vector. Negative control (intergenic), positive control (POLR2B), POMT1, and POMT2 guides were cloned into an mCherry-expressing vector. Luciferase-expressing, growth factor-independent BRAFV600E-expressing, or growth factor-independent KIAA1549::BRAF-expressing h9-NSCs were seeded into geltrex-coated 6-well plates at a density of 5×105 cells/well. Each well was supplemented with 15 mL of concentrated virus. After 16-24 hours the growth medium was changed to 2 mL of fresh TSM (luciferase-expressing cells) or TSM-EGF/bFGF (BRAFV600E- and KIAA1549::BRAF-expressing cells) per well. After 72 hours, cells were detached with accutase, and EGFP- and mCherry-expressing cells were pooled 1:1 as described in (Supplemental Table 5). The fraction of EGFP- and mCherry-expressing cells was analyzed on a Cytoflex flow cytometer. Cells were cultured for an additional 18 days, and the fraction of EGFP- and mCherry-expressing cells was assessed with flow cytometry.

### Immunoblotting

Cells were lysed in RIPA buffer supplemented with protease/phosphatase inhibitors according to the manufacturer’s protocol. Lysates were clarified at 20,000g for 10-20 minutes then NuPAGE sample buffer with 5% beta-mercaptoethanol was added and samples were boiled at 95C for 10 minutes. Samples were loaded onto a 4-12% bis-tris gradient gel and run at 150V for 2-3 hours. The gel was wet transferred to PVDF for 16-24 hours at 150 mA at room temperature. Membranes were blocked using the appropriate blocking solution described in (**Supplemental Table 8**). Membranes were incubated in primary antibody as described in (**Supplemental Table 8**) overnight for 16-24 hours at 4°C. The next day membranes were washed 3× in TBS-T for 5 minutes per wash. Membranes were incubated in the appropriate HRP-conjugated secondary antibody (**Supplemental Table 8**) for 1 hour at room temperature. Membranes were washed 3× in TBS-T for 5 minutes per wash. TBS-T was removed, and membranes were incubated with 1 mL of each SignalFire Elite (Cell Signaling Technology, #12757), SuperSignal West Pico PLUS (Thermo Fisher Scientific, #34580) or SuperSignal West Femto Maximum Sensitivity (Thermo Fisher Scientific, #34094) reagent for 1 minute with shaking before being imaged on the LI-COR Odyssey M.

### Immunofluorescence staining

Cells were plated onto geltrex-coated µ-Slide 8 Well chambered coverslip (Ibidi, #80807). After 24 hours of plating, cells were fixed with 3.7% paraformaldehyde for 15 minutes at room temperature. Cells were rinsed once with PBS then permeabilized with ice cold methanol for 15 minutes at 4°C. Cells were rinsed with PBS then blocked in 2% BSA diluted in PBS for one hour at room temperature. Cells were incubated in primary antibody (Supplemental Table 8) diluted in blocking buffer for 16-24 hours at 4°C. The next day cells were washed 3× in PBS (5-15 minutes per wash). Cells were incubated in secondary antibody (Supplemental Table 8) for 1 hour at room temperature, shielded from light. Cells were again washed 3× in PBS (5-15 minutes per wash) before being mounted in Ibidi Mounting Media with DAPI (Ibidi # NC1943852), cured at room temperature for 15 minutes, and stored shielded from light at 4°C prior to imaging.

### Confocal microscopy

Images taken for fusion protein localization analysis were acquired on a Zeiss LSM 980 microscope equipped with an Airyscan 2 detector (Carl Zeiss) using a LD LCI Plan-Apochromat 25x/0.8 Imm Corr DIC M27 objective. Bidirectional scanning with 2.05μseg pixel dwell time and averaging of 2 was used. Pixel size was set to 0.1657μm in compliance with Nyquist sampling and pixel intensities were encoded using 16-bit depth. Pinhole sizes were set to 0.8-1.4 Airy units. Excitation (detailing nominal laser power) and emission wavelengths were as follows: DAPI, *λ*_ex_ 405nm (at 15mV) - *λ*_em_ 408-501nm; Alexa Flour 488, *λ*_ex_ 488nm (at 13mV) - *λ*_em_ 491-588nm; Alexa Fluor 594, *λ*_ex_ 594nm (at 4mV) - *λ*_em_ 597-694nm. Airyscan post processing was performed using Zeiss ZEN software.

### Image analysis pipeline

Localization analysis of fusion proteins and the endoplasmic reticulum (labeled with PDI) was performed using CellProfiler 4.2.8111. Briefly, cells were segmented using the PDI channel and colocalization was assessed in the cytoplasmic region using Perason’s Correlation Coefficient. The code for all the analysis pipelines are available in the github repository associated with this manuscript (see Code Availability).

### Intracellular flow cytometry

Cells were detached with accutase, and 300,000 cells were seeded into each well of a round-bottom 96-well plate. Cells were pelleted at 300g for 3 minutes, then the supernatant was removed. Cells were washed 2x more in PBS. Cells were fixed in 3.7% paraformaldehyde for 15 minutes at room temperature and were washed 3× in PBS. Cells were permeabilized and blocked in 5% BSA-TBST + 0.1% Triton-X100 for 1 hour at room temperature. Cells were incubated primary antibody (**Supplemental Table 8**) for 2 hours at room temperature with agitation every 30 minutes. Cells were pelleted and washed 3× in PBS. Cells were incubated in PE-conjugated secondary antibody (**Supplemental Table 8**) for 1 hour at room temperature, with agitation every 30 minutes. Cells were washed 3× in PBS, then were resuspended in 200 mL of PBS + 2% BSA and were analyzed on a Cytoflex flow cytometer.

### Glycoproteomics

Sample preparation and analysis was as previously described47 with minor modifications. Briefly, mNSC cell pellets were extracted in 0.5% SDS, 40 mM Tris-HCl, pH 8.0, 150 mM NaCl before probe sonication for disruption of cellular debris and DNA. Each extract was reduced (10 mM DTT, 72°C, 15 min) and alkylated (25 mM IAA, RT, 15 min) before adding Triton X-100 to a final concentration of 1% (v/v). N-glycans were digested with 10U PNGase F enzyme (Roche), 37°C, 18h. N-deglycosylated protein extracts were subsequently acetone precipitated (−20°C, 16h) and resolubilized in 40 mM TrisHCl, pH 8.0, 150 mM NaCl buffer for digestion with 50 µg trypsin or 25 µg GluC at 37°C and 18h. Peptides were desalted on Sep-Pak C18 columns (50 mg) and labeled with diethyl stable isotopes47 before mixing the samples in 1:1 (v/v) ratio. C- and O-linked mannose glycopeptides were enriched in batch-mode with 200 µL agarose slurry conjugated to BC2L-A lectin, prepared as previously described47, in 1 mL 40 mM Tris-HCl, pH 8.0, 150 mM NaCl, 1 mM CaCl2 buffer. Following 3×2 mL washes of the agarose beads, the glycopeptides were eluted with 100 mL 40 mM Tris-HCl, pH 8.0, 150 mM NaCl, 1 mM CaCl2, 40 mM EDTA solution. Enriched glycopeptides were desalted on Stage-tip C18 columns and analyzed by mass spectrometry as previously described47. Data processing, identification and relative quantification of C-and O-Man glycopeptide abundances was with the Proteome Discoverer 1.4 software as previously described47.

### TAILS mass spectrometry

HEK-293T cells were transfected with pMIBerry KIAA1549::BRAF (16::9) plasmid and lysed in NLB 48 h later. 30 mL of anti-HA affinity matrix (Roche) was added to 900 µL of cleared lysate and incubated with end-over-end rotation at 4°C for 4 hours. Sample workup was performed as previously described^112^. Briefly, proteins were denatured with 6 M guanidine hydrochloride in HEPES (pH 7.8), reduced with 1 mM DTT, and alkylated with 5.5 mM IAA. N-terminal and lysine side chain amines were blocked by demethylation using formaldehyde and sodium cyanoborohydride. After acetone precipitation, proteins were resuspended and digested overnight with trypsin. Newly formed peptides with unblocked N-termini were bound overnight to HPG-ALDII polymer in the presence of sodium cyanoborohydride. Unbound blocked peptides were recovered with 30 kDa molecular weight cut-off spin filters. Filtrates were desalted with STAGE tips^113^. LC-MS/MS measurements were performed on an Exploris 480 mass spectrometer, coupled to an EasyLC 1200 nanoflow-HPLC (ThermoFisher Scientific). Peptides were separated on a fused silica HPLC column tip (I.D. 75 µm, New Objective, self-packed with ReproSil-Pur 120 C18-AQ, 1.9 µm to a length of 20 cm) using a gradient of A (0.1% formic acid in water) and B (0.1% formic acid in 80% acetonitrile in water). The mass spectrometer was operated in data-dependent mode; after each MS scan (mass range m/z = 370 – 1750; resolution: 120’000) a maximum of twenty MS/MS scans were performed using an isolation window of 1.3, a normalized collision energy of 28%, a target AGC of 50% and a resolution of 15’000. Spray voltage was set to 2.3 kV and the ion-transfer tube temperature to 250°C; no sheath and auxiliary gas were used.

Raw mass spectrometry files were analyzed using MaxQuant (version 2.0.1.0)^114^ using either a customized database containing the KIAA1549::BRAF (16::9) protein sequence and common contaminants, or together with a UniProt full-length human database with common contaminants including keratins and digestion enzymes using semispeific ArgC as enzyme specificity. Carbamidomethylcysteine and dimethylation of lysine were set as fixed modification and protein amino-terminal acetylation, oxidation of methionine and dimethylation of peptide N-termini were set as variable modifications. The MS/MS tolerance was set to 20 ppm. Peptide, site, and protein FDR based on a forward-reverse database were set to 0.01, and minimum number of peptides for identification of proteins was set to one, which must be unique. The “match-between-run” option was used with a time window of 0.7 min.

### Protease inhibitor screen

Growth factor-independent KIAA1549::BRAF-expressing h9-NSCs were seeded into geltrex-coated 6-well plates and were allowed to attach overnight. The next day cells were treated with 1 µM of each protease inhibitor available within a commercially available library (Enzo #BML-2833-0100, **Supplemental Table 7**), pomalidomide, or DMSO (vehicle control). Cells were treated for 48 hours before being lysed and immunoblotted as previously described.

### Animal studies

#### In Utero Electroporation Mouse Models

All IUE mouse work was conducted according to institutional guidelines and standards from the Institutional Animal Care and Use Committee review board (University of Cincinnati IACUC #22-05-15-01). CD1-ICR (Charles River code #022) mice were used for all IUE experiments. Briefly, IUE models were created by lateral ventricle injection of nucleic acid mixtures followed by electroporation as described previously (Patel et al, 2020 and Wei et al. 2021). All DNA plasmids were used at a final concentration of 1µg/ µL. KIAA1549::BRAF and BRAF^V600E^ plasmids were co-electroporated with PBCAG-EGFP and pCAG-PBase. Cas9 was used at a final concentration of 33.3µM. sgRNAs were used at a final concentration of 20µM.

#### Plasmid construction and CRISPR-Cas9 reagents

Piggybac DNA plasmids were constructed as previously described^115^. PBCAG-KIAA1549::BRAF plasmid were generated by PCR amplification of KIAA1549::BRAF or BRAF^V600E^ and insertion into linearized PBCAG-EGFP, which was digested with EcoR1 and Not1 to remove the EGFP coding sequence. Ligations were performed with InFusion Snap assembly (Takara) and plasmid sequence verified by whole plasmid sequencing (Plasmidsaurus). All plasmid stocks were prepared using NucleoBond Xtra Maxi or Midi EF endotoxin-free kits (Machery Nagel). For CRISPR/Cas9: Cas9 was purchased from Trilink (#L-7206-100). Positive control Rosa26 sgRNA and multi-guide Pomt1 gene KO kit (**Supplemental Table 10**) were purchased from Synthego (now Editco).

#### Tissue collection, processing and immunostaining

Following euthanasia by CO_2_ inhalation brains were dissected into ice cold Dulbecco’s phosphate-buffered saline (DPBS). Samples were drop fixed in freshly prepared 4% paraformaldehyde overnight at 4°C. The following day samples were washed in cold PBS and then transferred into a 30% sucrose solution for cryopreservation. Samples were embedded in optimal cutting temperature (OCT) embedding medium and stored at −80°C until sectioned. 45µm thick free-floating sections were cut on a cryostat (Leica) and stored in PBS + 0.05% sodium azide until processed for staining. For immunofluorescent staining, sections were first transferred to blocking solution (PBS + 0.5% Triton X-100 + 10% normal donkey serum) prior to addition of primary antibodies (**Supplemental Table 8**) and incubated at 4°C overnight. The next day sections were washed in PBS and then placed into blocking solution containing corresponding fluorescent secondary antibodies (**Supplemental Table 8**) for overnight incubation at 4°C. Finally, sections were counterstained with Hoechst (1:1,000) and washed in PBS prior to mounting on slides and coverslipping (Fisherbrand, Superfrost Plus slides, Thermo Fisher ProLong Gold Antifade Mountant; Fisherbrand Microscope Cover Glass 24x50). Slides were imaged using a Nikon A1 confocal microscope. Image processing and analysis were performed in NIH Fiji/ImageJ and Imaris.

#### Targeted sequencing for CRISPR/Cas9

Genomic DNA was harvested from mouse neural stem cell cultures generated from electroporated brains. Briefly, 24 hours post IUE, brains were harvested and dissociated into single cell suspensions by papain enzyme digestion. Neurosphere cell lines were established in NeuroCult media with mouse neural stem cell proliferation supplement and bFGF/EGF. Transfected cells were isolated by FACS sorting and gDNA was subsequently isolated from this purified transfected population. To amplify the Rosa locus the following PCR primers were used (**Supplemental Table 9**). PCR products were isolated by gel purification and submitted for Sanger sequencing at the CCHMC DNA sequencing core with the sequencing primer (**Supplemental Table 9**). To amplify and sequence the Pomt1 locus the following PCR primers were used (**Supplemental Table 9**).

## Notes

### Competing Interest Statement

PB has received grant funding from Novartis Institute of Bio-medical Research, and has served on paid advisory boards for QED Therapeutics, and Day One Biopharmaceuticals. RB consults for and owns equity in Scorpion Therapeutics, Kar-yoverse Therapeutics, and LOH Therapeutics. WCH is a consultant for Thermo Fischer, Solasta Ventures, KSQ Ther-apeutics, Frontier Medicines, Jubilant Therapeutics, RAPPTA Therapeutics, Serinus Biosciences, Kestral Therapeutics, Crane Biotherapeutics, Function Oncology, Recursion Pharma, and Weaver Biotherapetics and Perceptive. DER receives research funding from members of the
Functional Genomics Consortium (Abbvie, BMS, Jannsen, Merck, Vir), and
is a director of Addgene, Inc. TM is advisory board member for Ipsen Pharma GmbH. SAM is a consultant for Karyoverse Therapeutics and is a consultant for and owns equity in LOH Therapeutics. ESF is a founder, scientific advisory board (SAB) member, and equity holder of Civetta Therapeutics, Proximity Therapeutics, Neomorph, Inc. (also board of direc-tors), StelexisBiosciences, Inc., Anvia Therapeutics, Inc. (also board of directors) and CPD4, Inc. (also board of directors). He is an equity holder and SAB member for Avilar Therapeu-tics, PhotysTherapeutics, and Ajax Therapeutics and an equity holder in Lighthorse Therapeutics. E.S.F. is a consultant to Novartis, EcoR1 capital, Odyssey and Deerfield. The Fischer lab receives or has received research funding from Deerfield, Novartis, Ajax, Interline, Bayer and Astellas. TB received speaker honoraria from Pierre Fabre and the European Society for Medical Oncology (ESMO)

## References

1. O’Brien, S. G. et al. Imatinib compared with interferon and low-dose cytarabine for newly diagnosed chronic-phase chronic myeloid leukemia. N. Engl. J. Med. 348, 994–1004 (2003).

2. Gao, Q. et al. Driver fusions and their implications in the development and treatment of human cancers. Cell Rep. 23, 227–238.e3 (2018).

3. Sievert, A. J. et al. Duplication of 7q34 in pediatric low-grade astrocytomas detected by high-density single-nucleotide polymorphism-based genotype arrays results in a novel BRAF fusion gene. Brain Pathol. 19, 449–458 (2009).

4. Jones, D. T. W. et al. Oncogenic RAF1 rearrangement and a novel BRAF mutation as alternatives to KIAA1549:BRAF fusion in activating the MAPK pathway in pilocytic astrocytoma. Oncogene 28, 2119–2123 (2009).

5. Forshew, T. et al. Activation of the ERK/MAPK pathway: a signature genetic defect in posterior fossa pilocytic astrocytomas. J. Pathol. 218, 172–181 (2009).

6. Cin, H. et al. Oncogenic FAM131B-BRAF fusion resulting from 7q34 deletion comprises an alternative mechanism of MAPK pathway activation in pilocytic astrocytoma. Acta Neuropathol. 121, 763–774 (2011).

7. Jones, D. T. W. et al. Recurrent somatic alterations of FGFR1 and NTRK2 in pilocytic astrocytoma. Nat. Genet. 45, 927–932 (2013).

8. Shin, C. H., Grossmann, A. H., Holmen, S. L. & Robinson, J. P. The BRAF kinase domain promotes the development of gliomas in vivo. Genes Cancer 6, 9–18 (2015).

9. Ross, J. S. et al. The distribution of BRAF gene fusions in solid tumors and response to targeted therapy. Int. J. Cancer 138, 881–890 (2016).

10. Collins, V. P., Jones, D. T. W. & Giannini, C. Pilocytic astrocytoma: pathology, molecular mechanisms and markers. Acta Neuropathol. 129, 775–788 (2015).

11. Hanrahan, A. J., Chen, Z., Rosen, N. & Solit, D. B. BRAF - a tumour-agnostic drug target with lineage-specific dependencies. Nat. Rev. Clin. Oncol. 21, 224–247 (2024).

12. Usta, D. et al. A cell-based MAPK reporter assay reveals synergistic MAPK pathway activity suppression by MAPK inhibitor combination in BRAF-driven pediatric low-grade glioma cells. Mol. Cancer Ther. 19, 1736–1750 (2020).

13. Fangusaro, J. et al. A phase II trial of selumetinib in children with recurrent optic pathway and hypothalamic low-grade glioma without NF1: a Pediatric Brain Tumor Consortium study. Neuro. Oncol. 23, 1777–1788 (2021).

14. Kilburn, L. B. et al. The type II RAF inhibitor tovorafenib in relapsed/refractory pediatric low-grade glioma: the phase 2 FIREFLY-1 trial. Nat. Med. 30, 207–217 (2024).

15. Kim, P. et al. FusionGDB 2.0: fusion gene annotation updates aided by deep learning. Nucleic Acids Res. 50, D1221–D1230 (2022).

16. Chen, M. F. et al. Tumor-agnostic genomic and clinical analysis of BRAF fusions identifies actionable targets. Clin. Cancer Res. 30, 3812–3823 (2024).

17. Lavoie, H. & Therrien, M. Regulation of RAF protein kinases in ERK signalling. Nat. Rev. Mol. Cell Biol. 16, 281–298 (2015).

18. Park, E. et al. Cryo-EM structure of a RAS/RAF recruitment complex. Nat. Commun. 14, 4580 (2023).

19. Ghosh, S. et al. The cysteine-rich region of raf-1 kinase contains zinc, translocates to liposomes, and is adjacent to a segment that binds GTP-ras. J. Biol. Chem. 269, 10000–10007 (1994).

20. ICGC/TCGA Pan-Cancer Analysis of Whole Genomes Consortium. Pan-cancer analysis of whole genomes. Nature 578, 82–93 (2020).

21. Northcott, P. A. et al. Enhancer hijacking activates GFI1 family oncogenes in medulloblastoma. Nature 511, 428–434 (2014).

22. Selt, F. et al. Establishment and application of a novel patient-derived KIAA1549:BRAF-driven pediatric pilocytic astrocytoma model for preclinical drug testing. Oncotarget 8, 11460–11479 (2017).

23. Selt, F. et al. Generation of patient-derived pediatric pilocytic astrocytoma in-vitro models using SV40 large T: evaluation of a modeling workflow. J. Neurooncol. 165, 467–478 (2023).

24. Koduri, V. et al. Peptidic degron for IMiD-induced degradation of heterologous proteins. Proc. Natl. Acad. Sci. U. S. A. 116, 2539–2544 (2019).

25. Moriya, K. et al. Protein N-myristoylation is required for cellular morphological changes induced by two formin family proteins, FMNL2 and FMNL3. Biosci. Biotechnol. Biochem. 76, 1201–1209 (2012).

26. Sievert, A. J. et al. Paradoxical activation and RAF inhibitor resistance of BRAF protein kinase fusions characterizing pediatric astrocytomas. Proc. Natl. Acad. Sci. U. S. A. 110, 5957–5962 (2013).

27. Meinnel, T., Dian, C. & Giglione, C. Myristoylation, an ancient protein modification mirroring eukaryogenesis and evolution. Trends Biochem. Sci. 45, 619–632 (2020).

28. Terrell, E. M. & Morrison, D. K. Ras-mediated activation of the RAF family kinases. Cold Spring Harb. Perspect. Med. 9, a033746 (2019).

29. Park, E. et al. Architecture of autoinhibited and active BRAF-MEK1-14-3-3 complexes. Nature 575, 545–550 (2019).

30. Martinez Fiesco, J. A., Durrant, D. E., Morrison, D. K. & Zhang, P. Structural insights into the BRAF monomer-to-dimer transition mediated by RAS binding. Nat. Commun. 13, 486 (2022).

31. Rajakulendran, T., Sahmi, M., Lefrançois, M., Sicheri, F. & Therrien, M. A dimerization-dependent mechanism drives RAF catalytic activation. Nature 461, 542–545 (2009).

32. Khadka, P. et al. PPM1D mutations are oncogenic drivers of de novo diffuse midline glioma formation. Nat. Commun. 13, 604 (2022).

33. Abid, T. et al. Genome-wide pooled CRISPR screening in neurospheres. Nat. Protoc. 18, 2014–2031 (2023).

34. Morin, E. et al. A diverse landscape of FGFR alterations and co-mutations defines novel therapeutic strategies in pediatric low-grade gliomas. bioRxiv 2024.08.27.609922 (2024) doi:10.1101/2024.08.27.609922.

35. Manya, H. et al. Demonstration of mammalian protein O-mannosyltransferase activity: coexpression of POMT1 and POMT2 required for enzymatic activity. Proc. Natl. Acad. Sci. U. S. A. 101, 500–505 (2004).

36. Vester-Christensen, M. B. et al. Mining the O-mannose glycoproteome reveals cadherins as major O-mannosylated glycoproteins. Proc. Natl. Acad. Sci. U. S. A. 110, 21018–21023 (2013).

37. Povolo, L., Tian, W., Vakhrushev, S. Y. & Halim, A. Global view of domain-specific O-linked mannose glycosylation in glycoengineered cells. Mol. Cell. Proteomics 23, 100796 (2024).

38. Larsen, I. S. B. et al. Mammalian O-mannosylation of cadherins and plexins is independent of protein O-mannosyltransferases 1 and 2. J. Biol. Chem. 292, 11586–11598 (2017).

39. Reitman, Z. J. et al. Mitogenic and progenitor gene programmes in single pilocytic astrocytoma cells. Nat. Commun. 10, 3731 (2019).

40. GTEx Consortium. The GTEx Consortium atlas of genetic regulatory effects across human tissues. Science 369, 1318–1330 (2020).

41. Colussi, P. A., Taron, C. H., Mack, J. C. & Orlean, P. Human and Saccharomyces cerevisiae dolichol phosphate mannose synthases represent two classes of the enzyme, but both function in Schizosaccharomyces pombe. Proc. Natl. Acad. Sci. U. S. A. 94, 7873–7878 (1997).

42. Gandini, R., Reichenbach, T., Tan, T.-C. & Divne, C. Structural basis for dolichylphosphate mannose biosynthesis. Nat. Commun. 8, (2017).

43. Takahashi, T. Dolichyl-Phosphate Mannosyltransferase Polypeptide (DPM1-3). in Handbook of Glycosyltransferases and Related Genes 1637–1647 (Springer Japan, Tokyo, 2014).

44. Tsherniak, A. et al. Defining a Cancer Dependency Map. Cell 170, 564–576.e16 (2017).

45. Dempster, J. M. et al. Chronos: a cell population dynamics model of CRISPR experiments that improves inference of gene fitness effects. Genome Biol. 22, 343 (2021).

46. Bai, L., Kovach, A., You, Q., Kenny, A. & Li, H. Structure of the eukaryotic protein O-mannosyltransferase Pmt1-Pmt2 complex. Nat. Struct. Mol. Biol. 26, 704–711 (2019).

47. Orchard, M. G. et al. Rhodanine-3-acetic acid derivatives as inhibitors of fungal protein mannosyl transferase 1 (PMT1). Bioorg. Med. Chem. Lett. 14, 3975–3978 (2004).

48. Neubert, P. & Strahl, S. Protein O-mannosylation in the early secretory pathway. Curr. Opin. Cell Biol. 41, 100–108 (2016).

49. Klement, E. & Medzihradszky, K. F. Extracellular protein phosphorylation, the neglected side of the modification. Mol. Cell. Proteomics 16, 1–7 (2017).

50. Ochoa, D. et al. The functional landscape of the human phosphoproteome. Nat. Biotechnol. 38, 365–373 (2020).

51. Hornbeck, P. V. et al. PhosphoSitePlus, 2014: mutations, PTMs and recalibrations. Nucleic Acids Res. 43, D512–20 (2015).

52. Potel, C. M. et al. Uncovering protein glycosylation dynamics and heterogeneity using deep quantitative glycoprofiling (DQGlyco). Nat. Struct. Mol. Biol. (2025) doi:10.1038/s41594-025-01485-w.

53. Hallgren, J. et al. DeepTMHMM predicts alpha and beta transmembrane proteins using deep neural networks. bioRxiv 2022.04.08.487609 (2022) doi:10.1101/2022.04.08.487609.

54. Kyte, J. & Doolittle, R. F. A simple method for displaying the hydropathic character of a protein. J. Mol. Biol. 157, 105–132 (1982).

55. Varadi, M. et al. AlphaFold Protein Structure Database: massively expanding the structural coverage of protein-sequence space with high-accuracy models. Nucleic Acids Res. 50, D439–D444 (2022).

56. Jumper, J. et al. Highly accurate protein structure prediction with AlphaFold. Nature 596, 583–589 (2021).

57. Pei, J. & Grishin, N. V. Expansion of divergent SEA domains in cell surface proteins and nucleoporin 54. Protein Sci. 26, 617–630 (2017).

58. Diedrich, B. et al. Discrete cytosolic macromolecular BRAF complexes exhibit distinct activities and composition. EMBO J. 36, 646–663 (2017).

59. Bork, P. & Patthy, L. The SEA module: a new extracellular domain associated with O-glycosylation. Protein Sci. 4, 1421–1425 (1995).

60. Wreschner, D. H. et al. Generation of ligand-receptor alliances by “SEA” module-mediated cleavage of membrane-associated mucin proteins. Protein Sci. 11, 698–706 (2002).

61. Gordon, W. R. et al. Structural basis for autoinhibition of Notch. Nat. Struct. Mol. Biol. 14, 295–300 (2007).

62. Leon, K. et al. Structural basis for adhesion G protein-coupled receptor Gpr126 function. Nat. Commun. 11, 194 (2020).

63. Angliker, H., Wikstrom, P., Shaw, E., Brenner, C. & Fuller, R. S. The synthesis of inhibitors for processing proteinases and their action on the Kex2 proteinase of yeast. Biochem. J. 293 (Pt 1), 75–81 (1993).

64. Fugère, M. et al. Inhibitory potency and specificity of subtilase-like pro-protein convertase (SPC) prodomains. J. Biol. Chem. 277, 7648–7656 (2002).

65. Pritchard, C. A., Samuels, M. L., Bosch, E. & McMahon, M. Conditionally oncogenic forms of the A-Raf and B-Raf protein kinases display different biological and biochemical properties in NIH 3T3 cells. Mol. Cell. Biol. 15, 6430–6442 (1995).

66. Oncogenic BRAF Regulates B-Trcp Expression and NF-JB Activity in Human Melanoma Cells.

67. Woods, D. et al. Raf-induced proliferation or cell cycle arrest is determined by the level of Raf activity with arrest mediated by p21Cip1. Mol. Cell. Biol. 17, 5598–5611 (1997).

68. Mutations of Critical Amino Acids Aect the Biological and Biochemical Properties of Oncogenic A-Raf and Raf-1.

69. Oncogenic BRAF Regulates B-Trcp Expression and NF-JB Activity in Human Melanoma Cells.

70. Mutations of Critical Amino Acids Aect the Biological and Biochemical Properties of Oncogenic A-Raf and Raf-1.

71. Gronych, J. et al. An activated mutant BRAF kinase domain is sufficient to induce pilocytic astrocytoma in mice. J. Clin. Invest. 121, 1344–1348 (2011).

72. Farmer, H. et al. Targeting the DNA repair defect in BRCA mutant cells as a therapeutic strategy. Nature 434, 917–921 (2005).

73. Kryukov, G. V. et al. MTAP deletion confers enhanced dependency on the PRMT5 arginine methyltransferase in cancer cells. Science 351, 1214–1218 (2016).

74. Mavrakis, K. J. et al. Disordered methionine metabolism in MTAP/CDKN2A-deleted cancers leads to dependence on PRMT5. Science 351, 1208–1213 (2016).

75. Marjon, K. et al. MTAP deletions in cancer create vulnerability to targeting of the MAT2A/PRMT5/RIOK1 axis. Cell Rep. 15, 574–587 (2016).

76. Chan, E. M. et al. WRN helicase is a synthetic lethal target in microsatellite unstable cancers. Nature 568, 551–556 (2019).

77. van Wietmarschen, N. et al. Repeat expansions confer WRN dependence in microsatellite-unstable cancers. Nature 586, 292–298 (2020).

78. Cullen, P. J. Signaling mucins: the new kids on the MAPK block. Crit. Rev. Eukaryot. Gene Expr. 17, 241–257 (2007).

79. Cullen, P. J. Post-translational regulation of signaling mucins. Curr. Opin. Struct. Biol. 21, 590–596 (2011).

80. Carson, D. D. The cytoplasmic tail of MUC1: a very busy place. Sci. Signal. 1, e35 (2008).

81. Cullen, P. J. et al. A signaling mucin at the head of the Cdc42- and MAPK-dependent filamentous growth pathway in yeast. Genes Dev. 18, 1695–1708 (2004).

82. Yang, H.-Y., Tatebayashi, K., Yamamoto, K. & Saito, H. Glycosylation defects activate filamentous growth Kss1 MAPK and inhibit osmoregulatory Hog1 MAPK. EMBO J. 28, 1380–1391 (2009).

83. Bausewein, D., Engel, J., Jank, T., Schoedl, M. & Strahl, S. Functional similarities between the protein O-mannosyltransferases Pmt4 from bakers’ yeast and human POMT1. J. Biol. Chem. 291, 18006–18015 (2016).

84. Vadaie, N. et al. Cleavage of the signaling mucin Msb2 by the aspartyl protease Yps1 is required for MAPK activation in yeast. J. Cell Biol. 181, 1073–1081 (2008).

85. Yoshida-Moriguchi, T. & Campbell, K. P. Matriglycan: a novel polysaccharide that links dystroglycan to the basement membrane. Glycobiology 25, 702–713 (2015).

86. Akhavan, A., Crivelli, S. N., Singh, M., Lingappa, V. R. & Muschler, J. L. SEA domain proteolysis determines the functional composition of dystroglycan. FASEB J. 22, 612–621 (2008).

87. Vega, L. L. de la et al. Abstract 2857: Perspectives from a systematic review of kinase fusion oncogenes and access to corresponding kinase inhibitor clinical trials in pediatric, adolescent, and young adult solid malignancies. Cancer Res. 84, 2857–2857 (2024).

88. Milde, T. et al. Optimizing preclinical pediatric low-grade glioma models for meaningful clinical translation. Neuro. Oncol. 25, 1920–1931 (2023).

89. O’Hare, P. et al. Resistance, rebound, and recurrence regrowth patterns in pediatric low-grade glioma treated by MAPK inhibition: A modified Delphi approach to build international consensus-based definitions-International Pediatric Low-Grade Glioma Coalition. Neuro. Oncol. 26, 1357–1366 (2024).

90. Schenkein, D. Proteasome inhibitors in the treatment of B-cell malignancies. Clin. Lymphoma 3, 49–55 (2002).

91. Manya, H. et al. Protein O-mannosyltransferase activities in lymphoblasts from patients with alpha-dystroglycanopathies. Neuromuscul. Disord. 18, 45–51 (2008).

92. Haines, N., Seabrooke, S. & Stewart, B. A. Dystroglycan and protein O-mannosyltransferases 1 and 2 are required to maintain integrity of Drosophila larval muscles. Mol. Biol. Cell 18, 4721–4730 (2007).

93. Ichimiya, T. et al. The twisted abdomen phenotype of Drosophila POMT1 and POMT2 mutants coincides with their heterophilic protein O-mannosyltransferase activity. J. Biol. Chem. 279, 42638–42647 (2004).

94. Lommel, M., Willer, T. & Strahl, S. POMT2, a key enzyme in Walker-Warburg syndrome: somatic sPOMT2, but not testis-specific tPOMT2, is crucial for mannosyltransferase activity in vivo. Glycobiology 18, 615–625 (2008).

95. Lommel, M. et al. Correlation of enzyme activity and clinical phenotype in POMT1-associated dystroglycanopathies. Neurology 74, 157–164 (2010).

96. Beltrán-Valero de Bernabé, D. et al. Mutations in the O-mannosyltransferase gene POMT1 give rise to the severe neuronal migration disorder Walker-Warburg syndrome. Am. J. Hum. Genet. 71, 1033–1043 (2002).

97. Perez-Riverol, Y. et al. The PRIDE database at 20 years: 2025 update. Nucleic Acids Res. 53, D543–D553 (2025).

98. Hardin, E. C. et al. LOGGIC Core BioClinical Data Bank: Added clinical value of RNA-Seq in an international molecular diagnostic registry for pediatric low-grade glioma patients. Neuro. Oncol. 25, 2087–2097 (2023).

99. Duckert, P., Brunak, S. & Blom, N. Prediction of proprotein convertase cleavage sites. Protein Eng. Des. Sel. 17, 107–112 (2004).

100. Varadi, M. et al. AlphaFold Protein Structure Database in 2024: providing structure coverage for over 214 million protein sequences. Nucleic Acids Res. 52, D368–D375 (2024).

101. Ives, C. M. et al. Restoring protein glycosylation with GlycoShape. Nat. Methods 21, 2117–2127 (2024).

102. Benjamini, Y. & Hochberg, Y. Controlling the false discovery rate: A practical and powerful approach to multiple testing. J. R. Stat. Soc. 57, 289–300 (1995).

103. Kim, H. K. et al. Deep learning improves prediction of CRISPR-Cpf1 guide RNA activity. Nat. Biotechnol. 36, 239–241 (2018).

104. DeWeirdt, P. C. et al. Optimization of AsCas12a for combinatorial genetic screens in human cells. Nat. Biotechnol. 39, 94–104 (2021).

105. Doench, J. G. et al. Optimized sgRNA design to maximize activity and minimize off-target effects of CRISPR-Cas9. Nat. Biotechnol. 34, 184–191 (2016).

106. Sanson, K. R. et al. Optimized libraries for CRISPR-Cas9 genetic screens with multiple modalities. Nat. Commun. 9, 5416 (2018).

107. Bandopadhayay, P. et al. MYB-QKI rearrangements in angiocentric glioma drive tumorigenicity through a tripartite mechanism. Nat. Genet. 48, 273–282 (2016).

108. International Cancer Genome Consortium PedBrain Tumor Project. Recurrent MET fusion genes represent a drug target in pediatric glioblastoma. Nat. Med. 22, 1314–1320 (2016).

109. Weinberg, F. et al. Identification and characterization of a BRAF fusion oncoprotein with retained autoinhibitory domains. Oncogene 39, 814–832 (2020).

110. Röring, M. et al. Distinct requirement for an intact dimer interface in wild-type, V600E and kinase-dead B-Raf signalling. EMBO J. 31, 2629–2647 (2012).

111. Stirling, D. R. et al. CellProfiler 4: improvements in speed, utility and usability. BMC Bioinformatics 22, 433 (2021).

112. Kleifeld, O. et al. Identifying and quantifying proteolytic events and the natural N terminome by terminal amine isotopic labeling of substrates. Nat. Protoc. 6, 1578–1611 (2011).

113. Rappsilber, J., Mann, M. & Ishihama, Y. Protocol for micro-purification, enrichment, pre-fractionation and storage of peptides for proteomics using StageTips. Nat. Protoc. 2, 1896–1906 (2007).

114. Cox, J. & Mann, M. MaxQuant enables high peptide identification rates, individualized p.p.b.-range mass accuracies and proteome-wide protein quantification. Nat. Biotechnol. 26, 1367–1372 (2008).

115. Patel, S. K. et al. Generation of diffuse intrinsic pontine glioma mouse models by brainstem-targeted in utero electroporation. Neuro. Oncol. 22, 381–392 (2020).

