## Extended Data Figures 1-4 for "O-mannosylation and protein maturation check-points represent therapeutic opportunities in BRAF fusion protein oncogenesis"

**Extended Figure 1. Development and characterization of oncogene-dependent human neural stem cell models of pLGG. (A)** Schematic depicting the generation and maintenance of h9-hNSCs with expression of V5- and IKFZ3-degron tagged Luciferase (vector control), KIAA1549::BRAF, and $BRAF^{V600E}$. Neural stem cells transduced to express KIAA1549::BRAF or $BRAF^{V600E}$ were cultured without exogenous EGF/bFGF supplementation, rendering them oncogene dependent. **(B)** Schematic representation of IKFZ3 mediated protein degradation wherein pomalidomide facilitates the interaction between the IZKF3 degron tag and an E3 ubiquitin ligase[24,115], resulting in rapid and titratable degradation of tagged proteins. **(C)** Engineered cells were treated with the indicated concentrations of pomalidomide for 24 hours. Cells were fixed and stained with a V5 antibody and relative V5 expression was assessed with flow cytometry. Data are from three independent biological replicates. The x-axis represents relative fluorescence intensity, corresponding to expression levels of the V5/IZKF3 tagged protein, the y-axis depicts the relative density of cells with a given V5 expression level. Concentration of pomalidomide used is depicted by different colored curves. **(D)** Incucyte S3 growth curves of engineered h9-hNSC models grown -/+ exogenous EGF/bFGF. Cell growth was measured for a total of 7 days. Data are from three independent biological replicates. The y-axis depicts percent confluence, and the x-axis depicts time since initial seeding (in days). Error bars represent Standard Error of the Mean (SEM). Two-way ANOVA, with Sidak's multiple comparisons test (against luciferase control -EGF/bFGF), $p < 0.0001$****. **(E)** Incucyte S3 growth curves of engineered h9-hNSC models grown -/+ pomalidomide in the absence of exogenous EGF/bFGF. Cell growth was measured for a total of 7 days. Data are from three independent biological replicates. The y-axis depicts percent confluence, and the x-axis depicts time since initial seeding (in days). Error bars represent Standard Error of the Mean (SEM). Two-way ANOVA, with Sidak's multiple comparisons test (luciferase control -pom vs luciferase control + pom, mean diff = 5.7; KIAA1549::BRAF -pom vs KIAA1549::BRAF +pom, mean diff = -56; $BRAF^{V600E}$ -pom vs $BRAF^{V600E}$ +pom, mean diff = -53.8), $p \leq 0.0001$****. **(F)** Representative immunoblot of engineered h9-NSC models assessing levels of pERK, ERK, and Actin. Cells were grown in the absence of exogenous EGF/bFGF supplementation for 24 hours -/+ pomalidomide (1 μM) before being lysed and immunoblotted. Data are from three independent biological replicates. **(G)** Quantification of (f). One-way ANOVA, with Sidak's multiple comparisons test (vs luciferase -pom), $BRAF^{V600E}$ -Pom pval < 0.0001, K::B -pom pval < 0.0001, Luciferase +Pom pval = 0.8931, $BRAF^{V600E}$ +Pom pval = 0.1185, K::B +Pom pval = 0.7635.

### Extended Figure 1

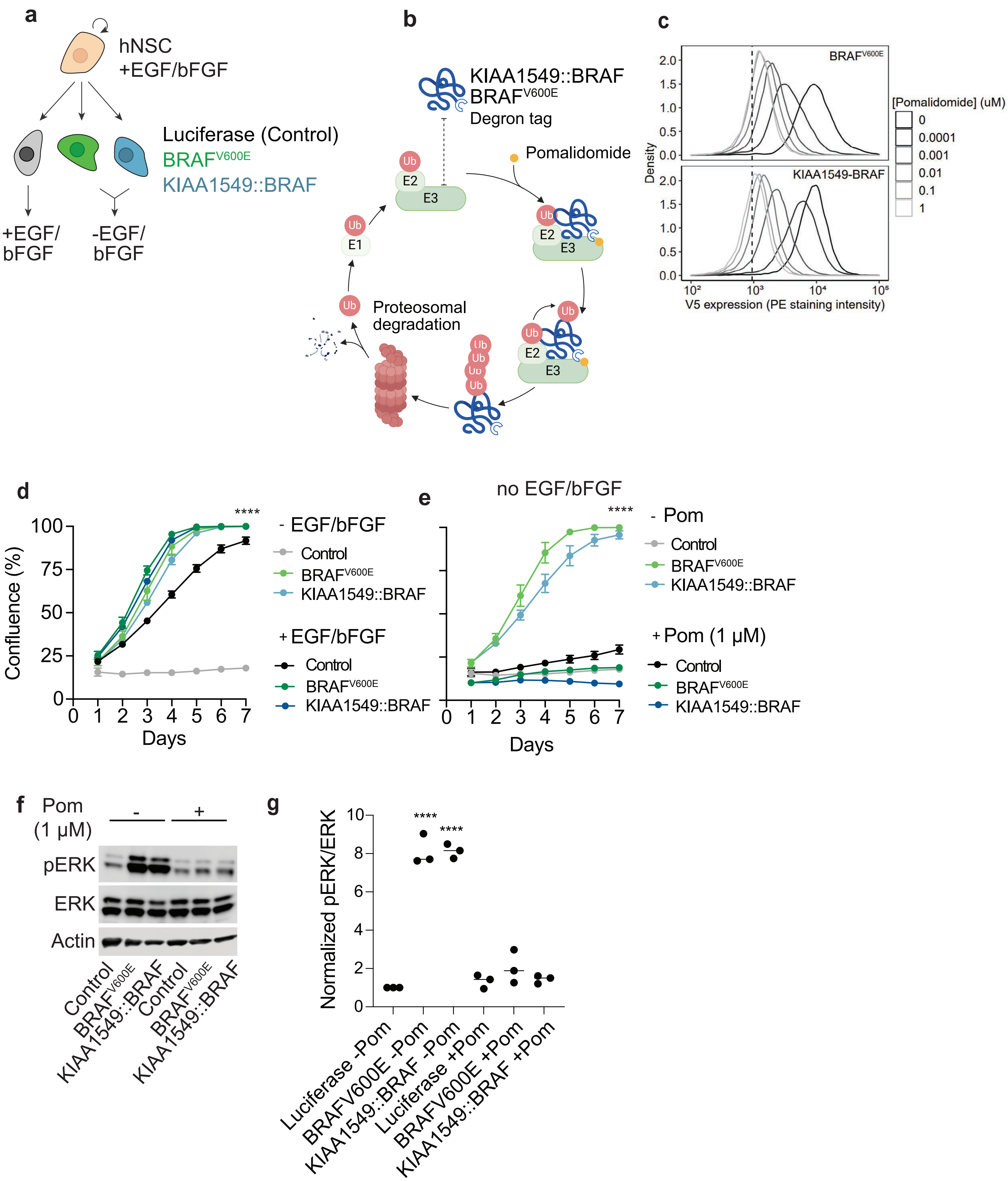

**Extended Figure 2. Development and characterization of oncogene-dependent human neural stem cell models of pLGG. (A)** Graphical representation of a genome-scale CRISPR/Cas9 screen in mNSC neurosphere lines. Isogenic mNSC models expressing Luciferase (control), $BRAF^{V600E}$, or KIAA1549::BRAF were infected with a genome-scale Cas9 library. Luciferase-expressing cells were grown with exogenous EGF/bFGF supplementation, and $BRAF^{V600E}$ and KIAA1549::BRAF-expressing cells were grown without. Following puromycin selection, cells were split and (early timepoint) genomic DNA samples were collected. Cells were cultured for an additional 18 days prior to (final timepoint) genomic DNA isolation. CRISPR/Cas9 guides were amplified with PCR and analyzed with next generation sequencing as described in Methods. **(B)** Bar graphs depicting relative expression levels of exogenously expressed human KIAA1549::BRAF and human $BRAF^{V600E}$ in mNSCs. Average expression across three independent biological replicates in presented as $\log_2$ fold change relative to luciferase-expressing mNSCs, normalized to B2M mRNA expression levels. Error bars represent the standard error of the mean. Student's t-test was used to compute statistical significance. Expression of K::B 15::9 breakpoint in K::B vs $BRAF^{V600E}$: p = 0.001**. Expression of BRAF exons 3-4 $BRAF^{V600E}$ vs K::B: p = 0.004**. **(C)** Incucyte growth curves demonstrating growth rates of engineered, oncogene-dependent, mNSC cell lines. Cells were engineered to express Luciferase, $BRAF^{V600E}$, or KIAA1549::BRAF and growth rates were measured for a total of 8 days in the presence (left) or absence (right) of exogenous EGF/bFGF supplementation. Data are from three independent biological replicates. Two-way ANOVA, with Sidak's multiple comparisons test: (against Control -EGF/bFGF), luciferase control + EGF/bFGF p < 0.0001****; $BRAF^{V600E}$ +/-EGF/bFGF p < 0.0001****; K::B + EGF/bFGF p < 0.0001****; K::B -EGF/bFGF p < 0.0009***. **(D)** Representative neurosphere images from **(C)**.

**Extended Figure 2**

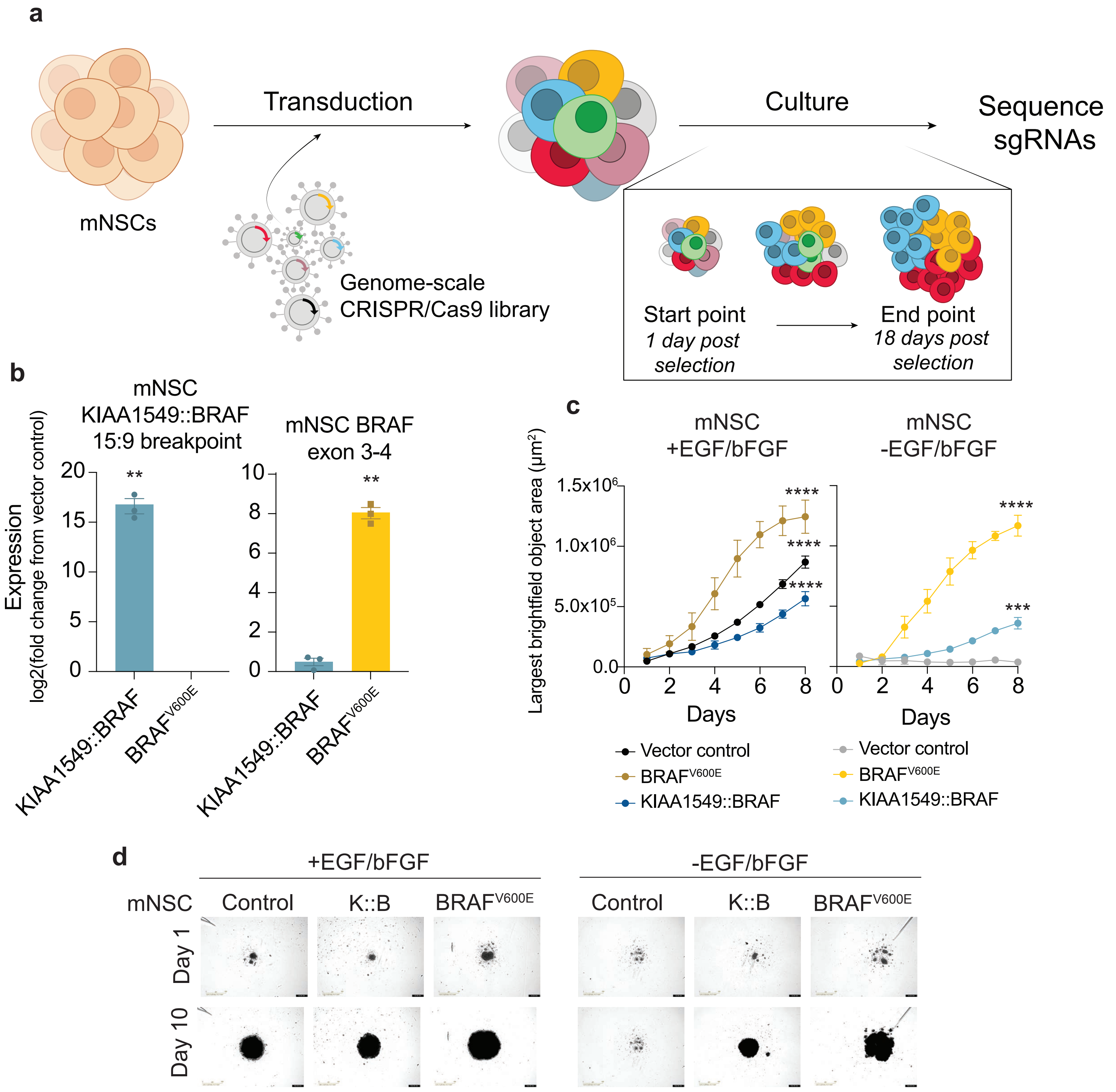

**Extended Figure 3. Expression of POMT1/2 and DPM1-3 in pLGG tumor samples.** Bulk RNA-Seq data from pLGG tumor samples (described in **Supplemental 36)**. **(A)** Association between POMT1/2 and DPM1-3 expression and driver oncogene status. Samples were stratified into those with a KIAA1549::BRAF fusion, a $BRAF^{V600E}$ point mutation, or those where the driver oncogene is not BRAF or is undetermined. Points represent individual tumor samples. The x-axis depicts the driver oncogene and the y-axis depicts the $log_2(TPM+1)$ mRNA expression for the indicated gene. Statistically significant differences in expression (relative to KIAA1549::BRAF-expressing tumors) were computed with Student's t-test with Bonferroni correction ($p < 0.05$) and are indicated with a *. **(B)** Correlation between POMT1/2 and DPM1-3 expression (as described in A) and patient age at time of sample collection. Samples are restricted to include only those with a KIAA1549::BRAF rearrangement. Pearsons's correlation coefficient was computed. Both p-values and $r^2$ values are indicated on each plot.

**Extended Figure 3**

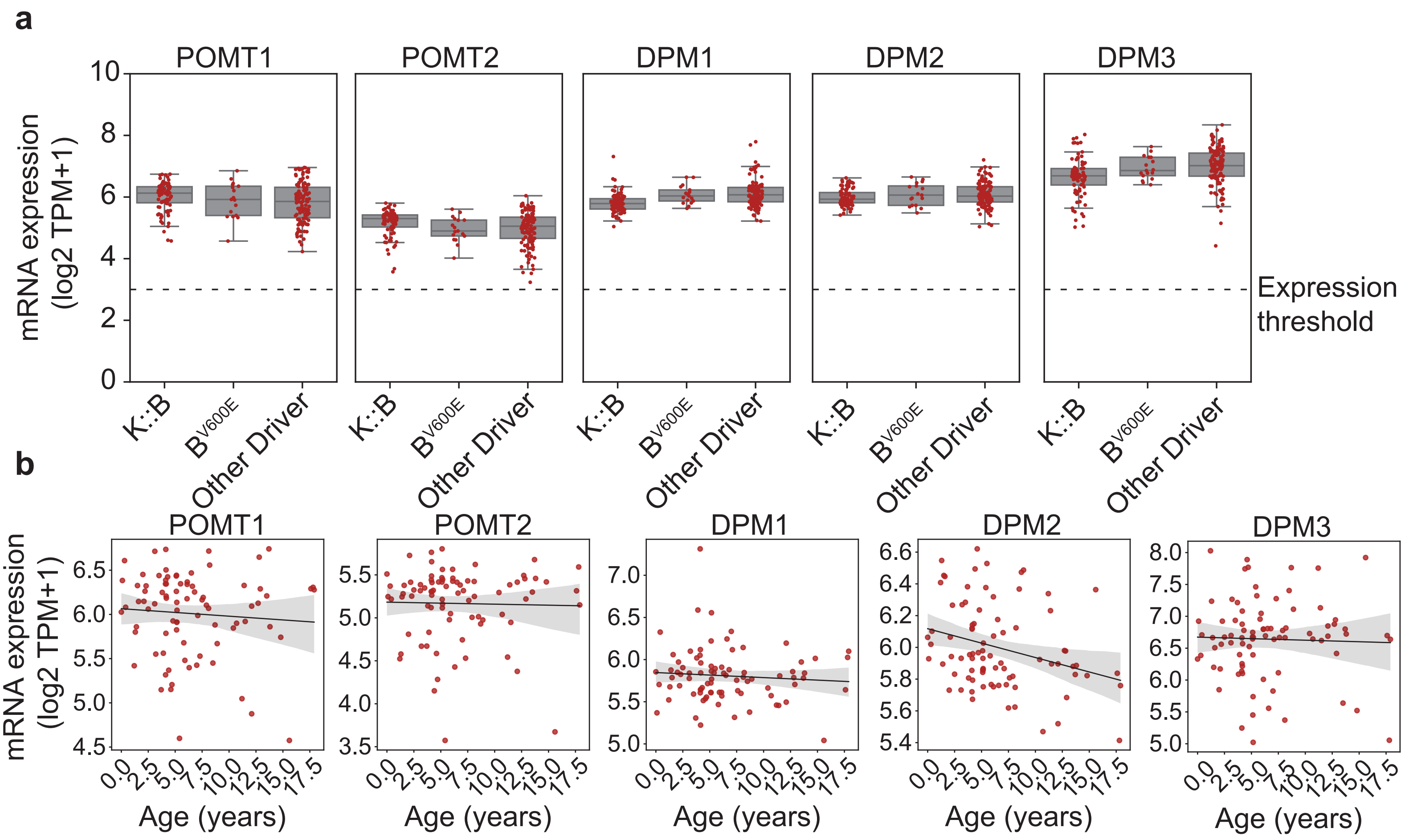

**Extended Figure 4. AlphaFold 2 predicted structure of putative juxtamembrane folded domains of KIAA1549.** Predicted juxtamembrane region domains and transmembrane region on KIAA1549. The predicted protein structure from residues 942-1324 of KIAA1549, including the folded domains (blue, domains 1-3) with paucimannose N-glycans grafted on (standard 3D-SNFG notation and colors), and the predicted transmembrane domain (red). The topology of the protein is indicated in the membrane, with the folded domains and N-linked glycosylation sites in the extracellular space. N-glycan structures and orientations are only representative, and do not necessarily represent the true glycan structures and conformations on KIAA1549

**Extended Figure 4**

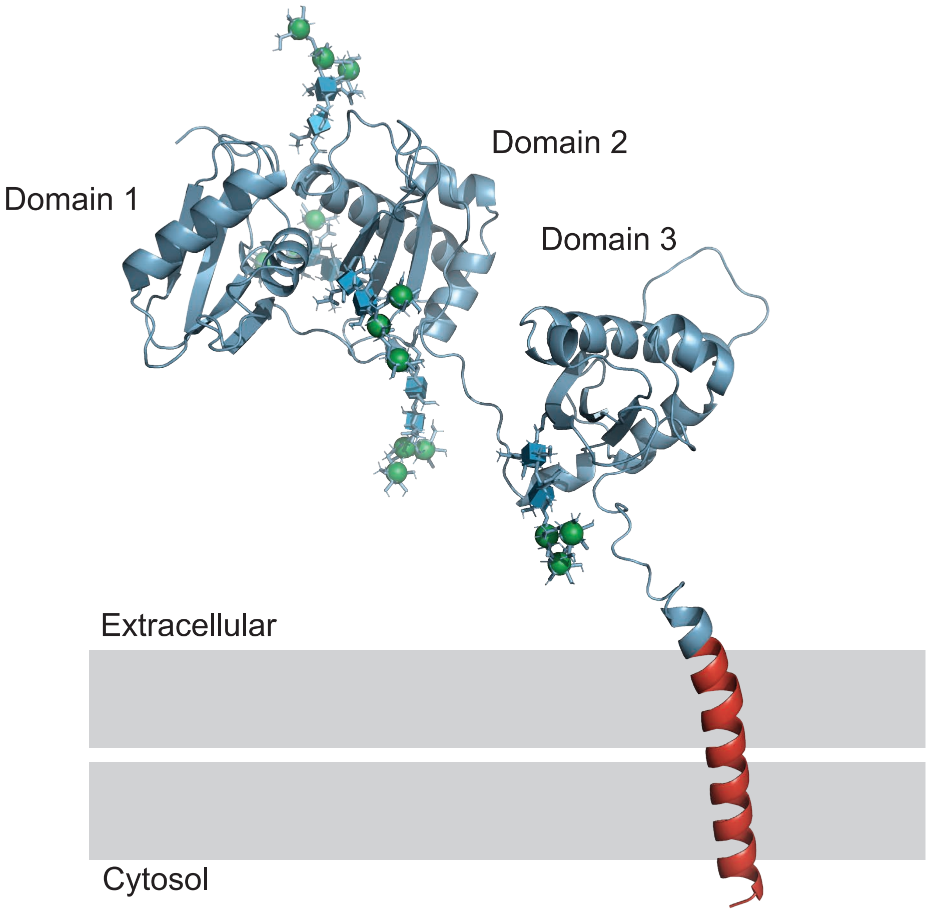
