## Supplemental Data Figures 1-9 for "O-mannosylation and protein maturation check-points represent therapeutic opportunities in BRAF fusion protein oncogenesis"

**Supplemental Figure 1.** (**A**) The number of unique fusions and the corresponding proto-oncogene for fusions in FusionGDB[15]. (**B**) Quantification of immunoblot in Fig. 1G, ERK and phospho-ERK in Luciferase (control), FAM131B::BRAF and FAM131B::BRAF$^{G2A}$. The level of phosphorylated ERK was normalized to total ERK. The plot shows mean value across all replicates (horizontal bar) and dots represent each biological replicate (n=3). Statistical analysis was performed with two-tailed ANOVA with Sidak's multiple hypothesis testing. Values were normalized to signal intensity of FAM131B::BRAF. Luciferase $p < 0.001$, FAM131B::BRAF$^{G2A}$ pval $< 0.001$. **(C)** Representative Airyscan images of h9-hNSCs expressing V5-tagged FAM131B::BRAF and FAM131B::BRAF$^{G2A}$. FAM131B::BRAF expression was detected with an antibody against the C-terminal V5 tag.

Supplemental Figure 1

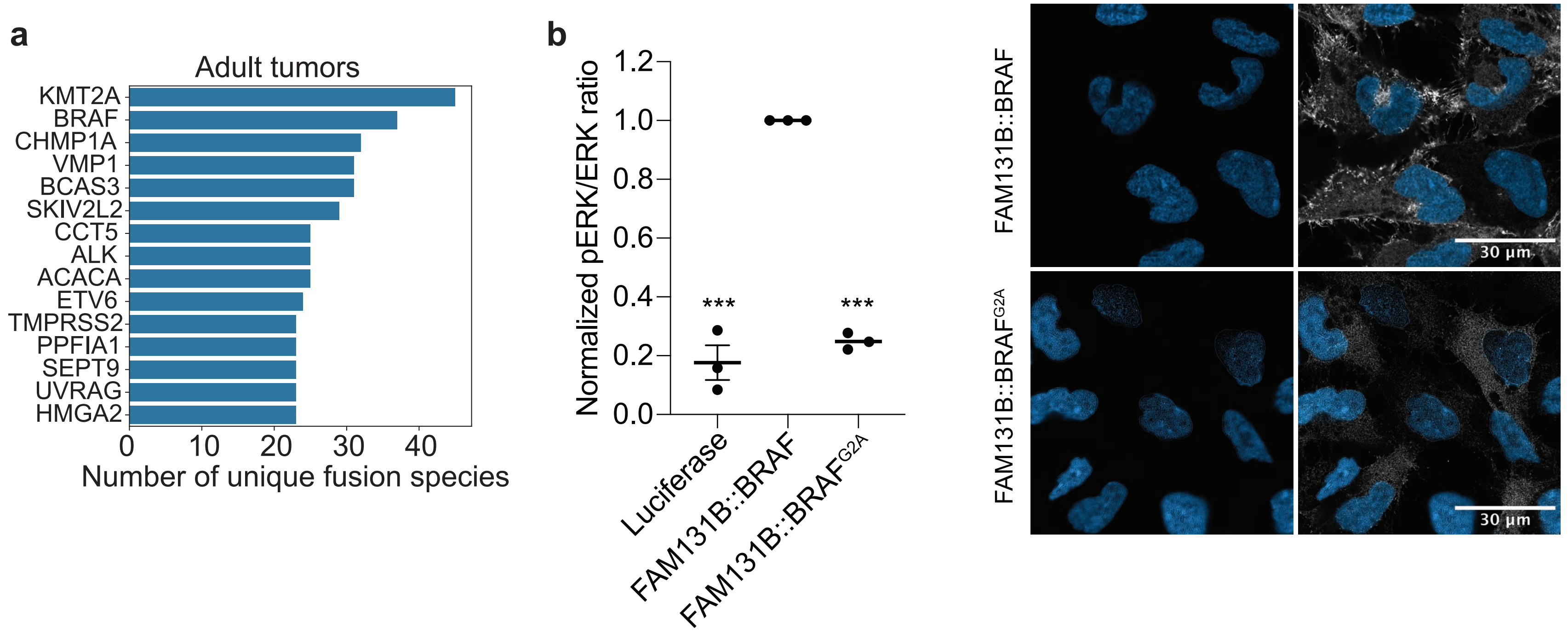

**Supplemental Figure 2.** Representative UMAPs (uniform manifold approximations and projections) were obtained using unsupervised clustering in Seurat, colored by their cell type annotation. Individual cells in the UMAP embedding for potentially malignant (Astrocyte, OPC, Oligodendrocyte, Progenitor Glia) or potentially non-malignant (Endothelial, Myeloid, Neuron, Stromal, T-cell) clusters, colored by expression of *POMT1*, *POMT2*, *DPM1*, *DPM2*, *DPM3*. Expression values are normalized for quantitative comparison across UMAP projections.

**Supplemental Figure 2**

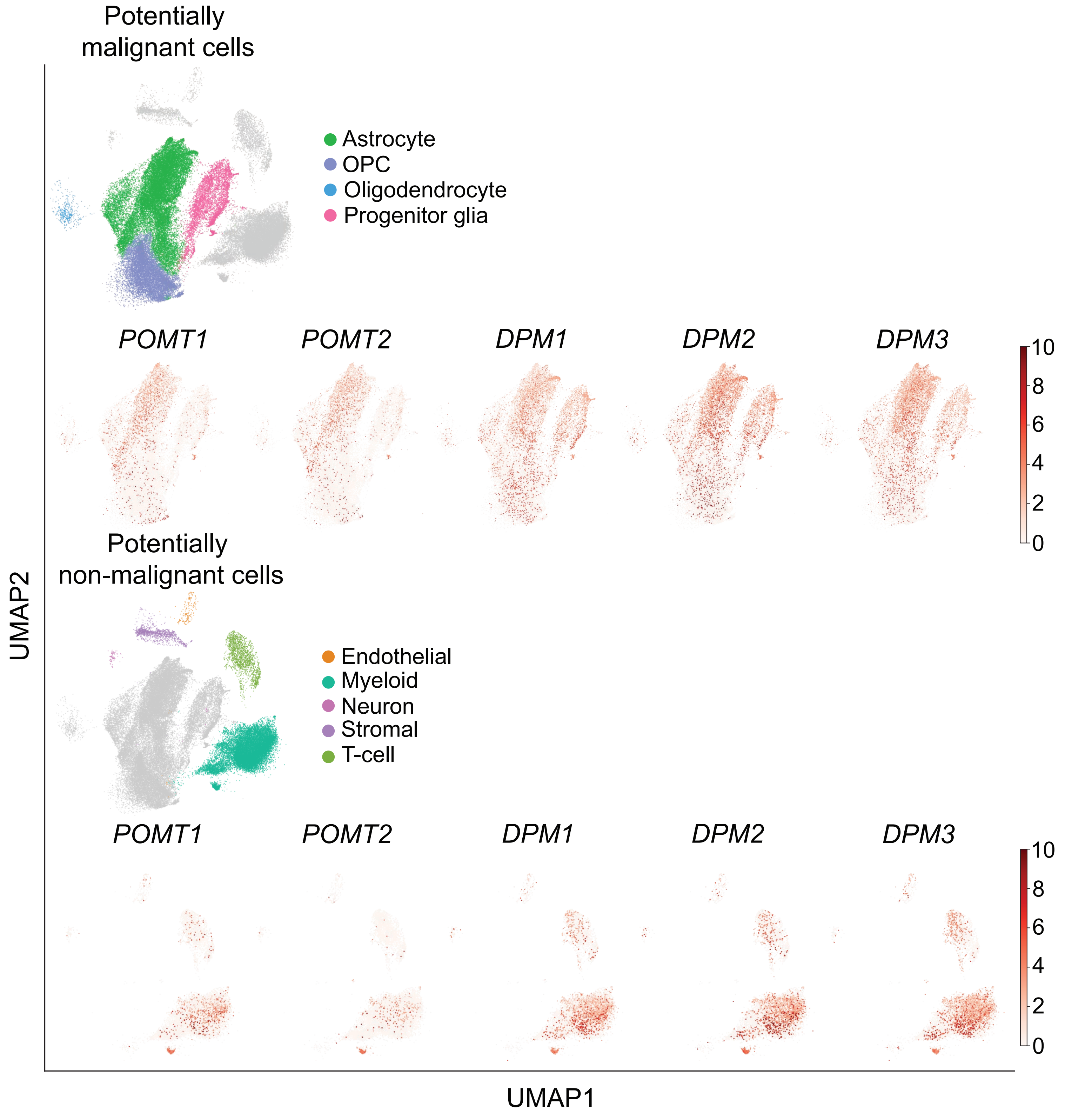

**Supplemental Figure 3.** Expression levels (TPM) of *POMT1/2* and *DPM1-3* from normal human tissue. Yellow bars represent cerebellum and the cerebellar hemisphere. The data and graphs were obtained from the GTEx Portal on 12/13/2024. The Genotype-Tissue Expression (GTEx) Project was supported by the Common Fund of the Office of the Director of the National Institutes of Health, and by NCI, NHGRI, NHLBI, NIDA, NIMH, and NINDS.

**Supplemental Figure 3**

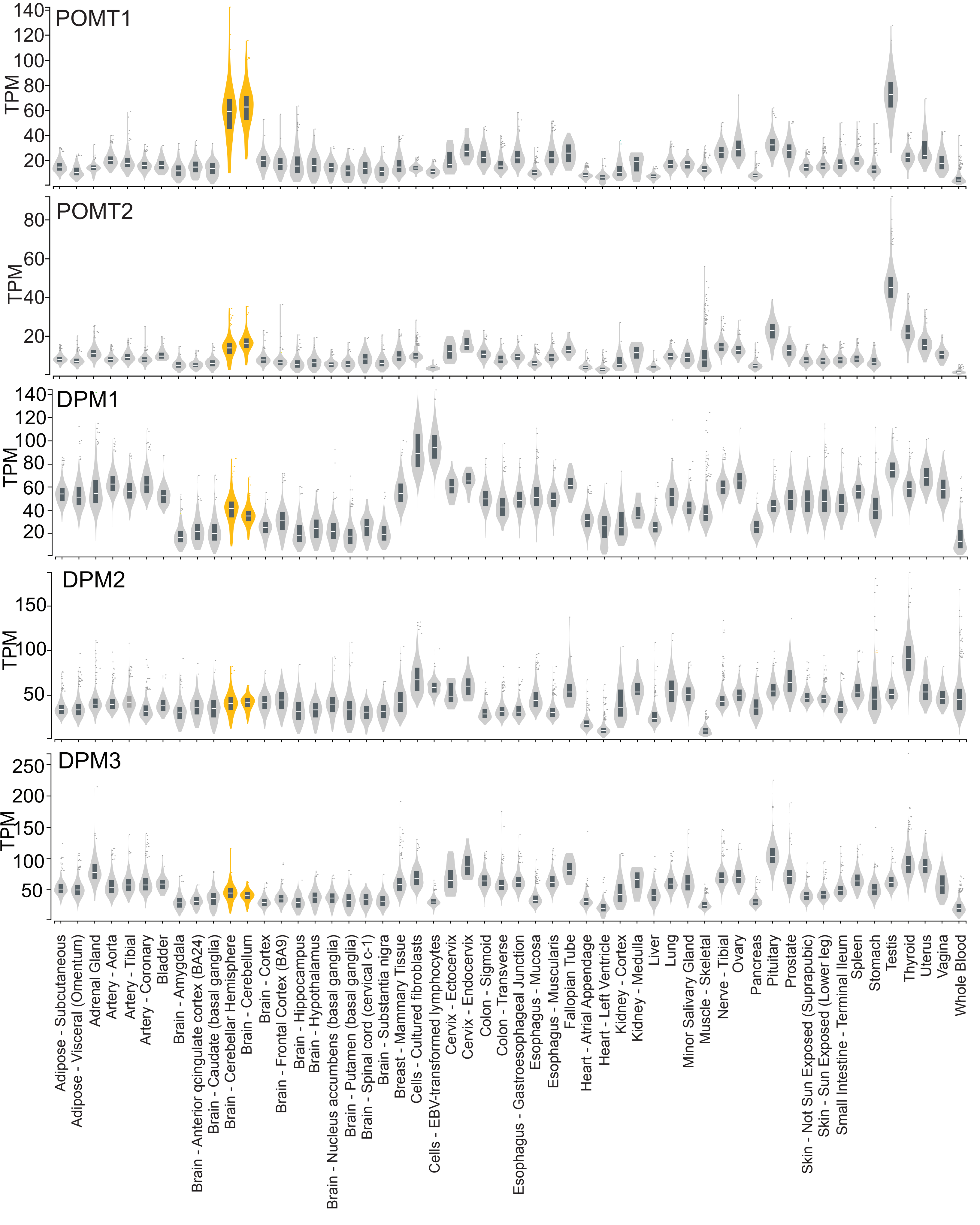

**Supplemental Figure 4. (A)** Cumulative cell doublings of mNSCs with expression of Luciferase (Control) or KIAA1549::BRAF (K::B) over 18 days post transduction of all-in-one vector carrying Cas9 and a guide against *Pomt1*, *Pomt2* or GFP (sgControl). Two different guides against *Pomt1* and *Pomt2* were used. Each isogenic cell model was cultured under their normal +/- EGF/bFGF condition as stated in the Figure. Cumulative doublings were calculated as change from day 1 post selection. Error bars represent the standard error (SEM) over three experimental replicates (n=3) and statistical differences between guide conditions and doublings over time were evaluated with t-tests with Welch's correction, sgGFP vs sg*Pomt1*, p = 0.007**. sgGFP vs sg*Pomt2*, p = 0.02*. There was no significant difference between guide conditions in the luciferase control mNSCs. **(B)** Bar graphs depicting percentage of frameshift, in-frame and uncut reads from CRISPRseq (n=1) of conditions in (b) at day 4, day 6 and Day 18 post viral transduction of all-in-one CRISPR/Cas9 vector. **(C)** Chronos score over all cell lines in DepMap for the pan-essential gene *POLR2B* (positive control for CRISPR competition assay in Fig. 3D). A gene with chronos score <-1 is considered essential/dependency. **(D)** Best-fit curve of mNSC isogenic models expressing Luciferase (control), KIAA1549::BRAF, KIAA1549 wild-type (WT), $BRAF^{V600E}$ and, BRAF wild-type (WT) treated with increasing concentrations of the POMT complex inhibitor R3A-5a and viability measured with CellTiterGlo after 72 hours of treatment. Cell viability was normalized to each equivalent DMSO control. Viability was assessed using area under the curve (AUC) of each experimental replicate (n=3) and then compared each cell line with luciferase control using one-way ANOVA with Sidak's multiple comparisons test: luciferase vs K::B p=0.0018**; luciferase vs KIAA1549 WT/ $BRAF^{V600E}$/BRAF WT p>0.9. (AUC luciferase control: 220.8; KIAA1549::BRAF: 140.8; KIAA1549 WT:227.7; $BRAF^{V600E}$ : 233; BRAF WT: 216.2).

#### Supplemental Figure 4

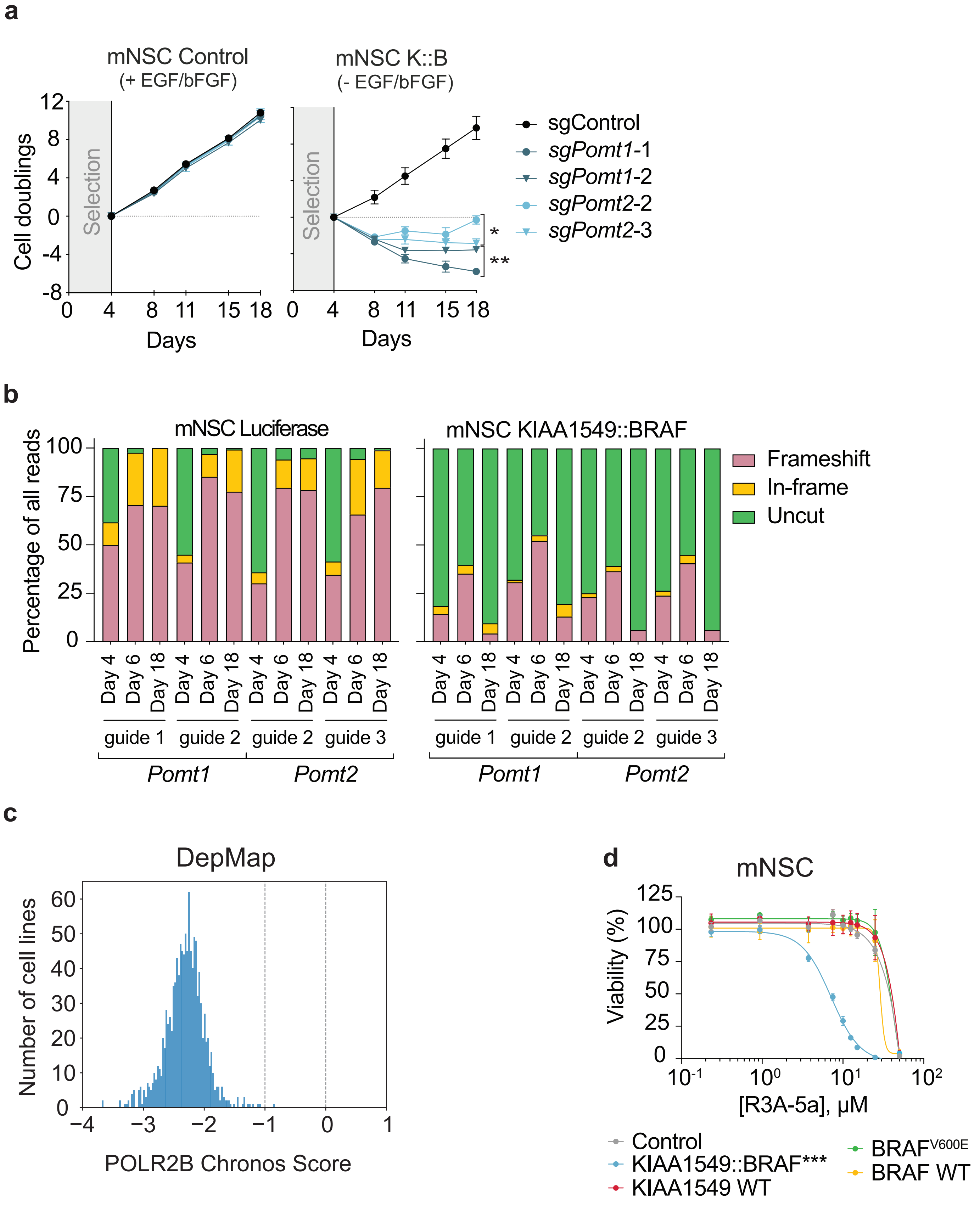

**Supplemental Figure 5. (A)** Immunoblot of lysates from $HEK293^{SC}$ with expression of K::B, treated with PNGaseF and/or POMT KO. **(B)** Proteomic analysis of mNSC VC (light) and mNSC K::B (heavy) cell digests. The tryptic digests were mixed in 1:1 (v/v) ratio and the mixing ratio was quantified at the peptide-level to estimate biological/technical variability. The frequency plot shows counts (n=2463) for log10 transformed heavy-to-light ratios of peptide pairs identified by the proteomic analysis. Interquartile range (IQR) calculations of Log10(H/L) ratios determined the boundaries ($-0.58 \leq x \leq 0.44$) for the technical/biological variability in the assay.

**Supplemental Figure 5**

**a**

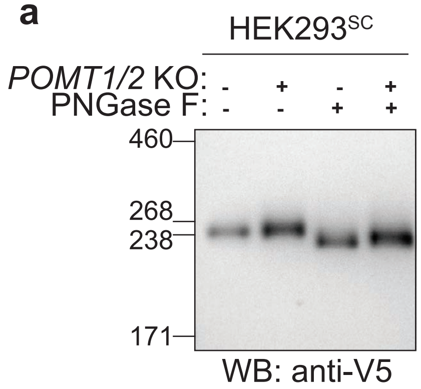


**b**

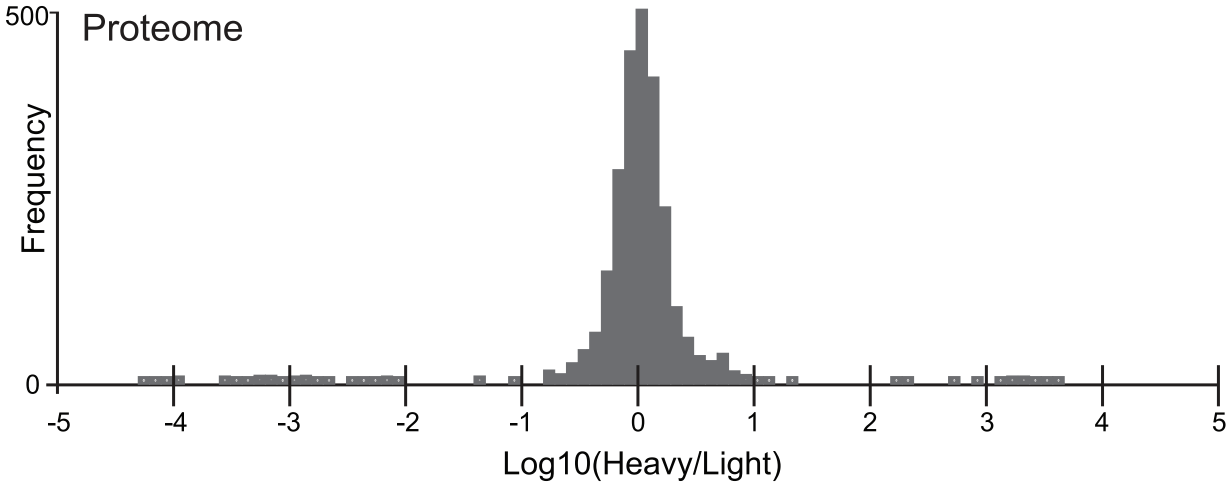

**Supplemental Figure 6.** Quantification of immunoblot in Fig. 2E, ERK and phospho-ERK in Luciferase (control), KIAA1549::BRAF and KIAA1549::BRAF$^{delTM}$ (deleted transmembrane domain). The level of phosphorylated ERK was normalized to total ERK. The plot shows mean value across all replicates (horizontal bar) and, dots represent each biological replicate (n=3). Statistical analysis was performed with two-tailed ANOVA with Sidak's multiple hypothesis testing. Values were normalized to signal intensity in the luciferase control condition. KIAA1549::BRAF pval < 0.05, KIAA1549::BRAF$^{delTM}$ pval > 0.05.

**Supplemental Figure 6**

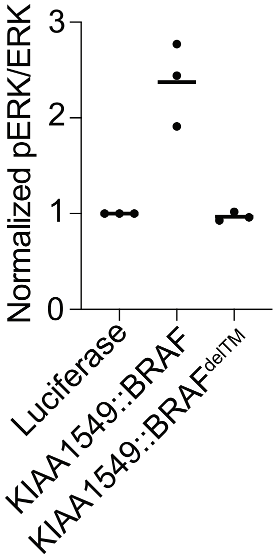

**Supplemental Figure 7.** (**A**) Quantification of the immunoblots in Fig 5a. The band intense for cleaved (110 kDa) and uncleaved (~240 kDa) K::B was quantified and the fraction of K::B in the cleaved state was plotted as the fraction: cleaved/(cleaved+uncleaved). Statistical analysis was performed with a Student's t-test, $p < 0.0001$. (**B**) Western blot analyses of HEK293T, mouse embryonic fibroblasts (MEF) and normal human astrocytes (NHA) transfected with BRAF wildtype and V600E mutant and three KIAA1549::BRAF breakpoint variants. Cells were transfected, lysed 48h later and subjected to Western blot analysis using the indicated antibodies. (**C**) Quantification of uncleaved (full length) KIAA1549::BRAF in immunoblot Fig. 5F. Graph shows each replicate (n=3) and the mean fraction of K::B in the uncleaved state. (**D**) Immunoblot of HA-tagged wildtype KIAA1549 and KIAA1549::BRAF in HEK293Ts. HEK293Ts were also engineered to express HA-tagged, mutated variants of KIAA1549 and KIAA1549::BRAF in which cleavage residues R1211 and R1212 were mutated to alanine. (**E**) Immunoblot showing full-length (top bands) KIAA1549 wild-type (WT) and proposed cleavage product (bottom bands) in fractionated lysates from HEK293Ts expressing HA-tagged KIAA1549 (WT). Blots showing whole-cell lysate (WCL), cytoplasmic, and membrane fractions with controls GAPDH and EGFR. (**F**) Full length K::B and the proposed cleavage product in immunolbots against the V5-tag on K::B in h9-hNSCs. Lysates from a protease screen of 51 protease inhibitors were analyzed (Furin Inhibitor 1 fold change vs DMSO = 11.5). (**G**) Incucyte growth curve from replicate one of three independent biological replicates. H9-NSCs were transduced with Luciferase, KIAA1549::BRAF, or KIAA1549::BRAF$^{delRMWR}$ and cells were grown for 6.75 days in the *presence* of exogenous EGF and bFGF supplementation. The x-axis depicts the time since the initial image, and the y-axis represents the relative cell confluence (normalized to confluence at 0h). Error bars represent Standard Error of the Mean (SEM). Statistics were calculated with two-way ANOVA across three biological replicates, Tukey's multiple comparison test (Control vs K::B, Control vs K::B$^{delRMWR}$ and K::B vs K::B$^{delRMWR}$): $p = 0.78$, $p = 0.18$, $p = 0.09$. (**H**) Incucyte growth curve from replicate two and three of three independent biological replicates. H9-NSCs were transduced with Luciferase (Control), KIAA1549::BRAF, or KIAA1549::BRAF$^{delRMWR}$ and cells were grown for 6.75 days in the *presence or absence* of exogenous EGF and bFGF supplementation. The x-axis depicts the time since the initial image, and the y-axis represents the relative cell confluence (normalized to confluence at 0h). Error bars represent Standard Error of the Mean (SEM). Statistics were calculated with two-way ANOVA across all three biological replicates, Tukey's multiple comparison test (Control vs K::B, Control vs K::B$^{delRMWR}$ and K::B vs K::B$^{delRMWR}$ in the *presence or absence* of EGF/bFGF): -EGF/bFGF: $p < 0.0001$****; +EGF/bFGF: $p = 0.78$, $p = 0.18$, $p = 0.09$. (same as H). (**I**) Quantification of immunoblots showing normalized ERK/pERK ratio in h9-hNSCs expressing luciferase control, K::B and K::B$^{delRMWR}$. Ratios were normalized to the pERK/ERK ratio in the luciferase control. Statistical analysis was performed with One-way ANOVA and with multiple hypothesis correction with Sidak's test. A p-value < 0.05* was considered statistically significant. (**J**) Plots represent the ratio of Band 2 (lower molecular weight band) to Band 1 (full-length form) across three biological replicates. Ratios are shown as mean ± Standard Error of the Mean (SEM) for each condition: WT, COSMC/POMGNT1 KO, and COSMC/POMGNT1/POMT1-2 KO. Statistical significance was evaluated using an unpaired t-test (Band 2/Band 1) (WT vs COSMC/POMGNT1 KO: $p = 0.087$. WT vs COSMC/POMGNT1/POMT1-2 KO: $p = 0.042$*).

### Supplemental Figure 7

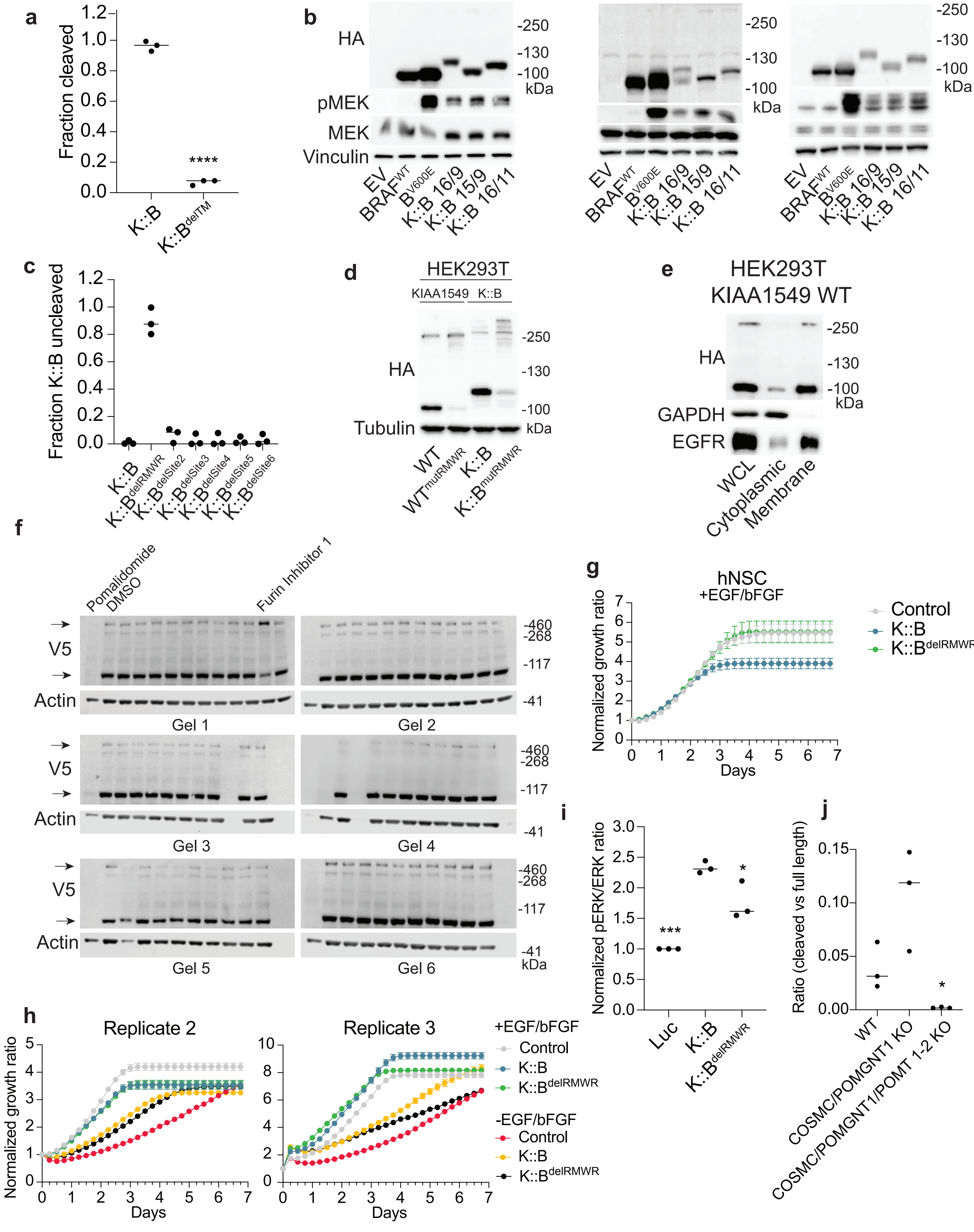

**Supplemental Figure 8**. **(A)** Close-up of domain 3 on KIAA1549. The two cleavage sites are indicated (orange stick representation), and the β2/β3 loop where cleavage is typically found on SEA domains is also indicated. **(B)** SEA domain gallery. Renderings of putative SEA domains from KIAA1549 and known SEA domains. The structures are colored following a rainbow color scheme from N- (blue) to C-terminal (red). The three KIAA1549 domains, and the Dystroglycan domain are extracted from Alphafold 2 predictions. The remaining domains are experimentally determined structures: Nup54 (from Xenopus Laevis, PDB 5C2U) and Muc16 (from Mouse, PDB 1IVZ). **(C)** Clustal Omega alignment of the RXXR motif cleavage site on KIAA1549 in orthologous proteins. An extract of the full sequence alignment for orthologous proteins (identified using matches for PFAM family) of KIAA1549. Representative organisms across metazoa were selected: Placozoa -Trichoplax adaherens (UniProt: B3RLT3), Cnidaria - Hydra (UniProt: A0A8B7DCG9), Anthozoa – Staghorn coral (UniProt: A0AAD9QQU3), Common octopus (UniProt: A0A7E6ESG4), Florida lancelet (UniProt: A0A9J7MND0), Lamprey (UniProt: A0AAJ7STN5). The motif for autoproteolytic cleavage on SEA domains is marked in green, while the Furin (RXXR) cleavage motif is marked in pink. N-linked sequons are marked in blue. Start and end residues for each protein are indicated, and the conservation of residues is indicated at the top using Clustal shading.

#### Supplemental Figure 8

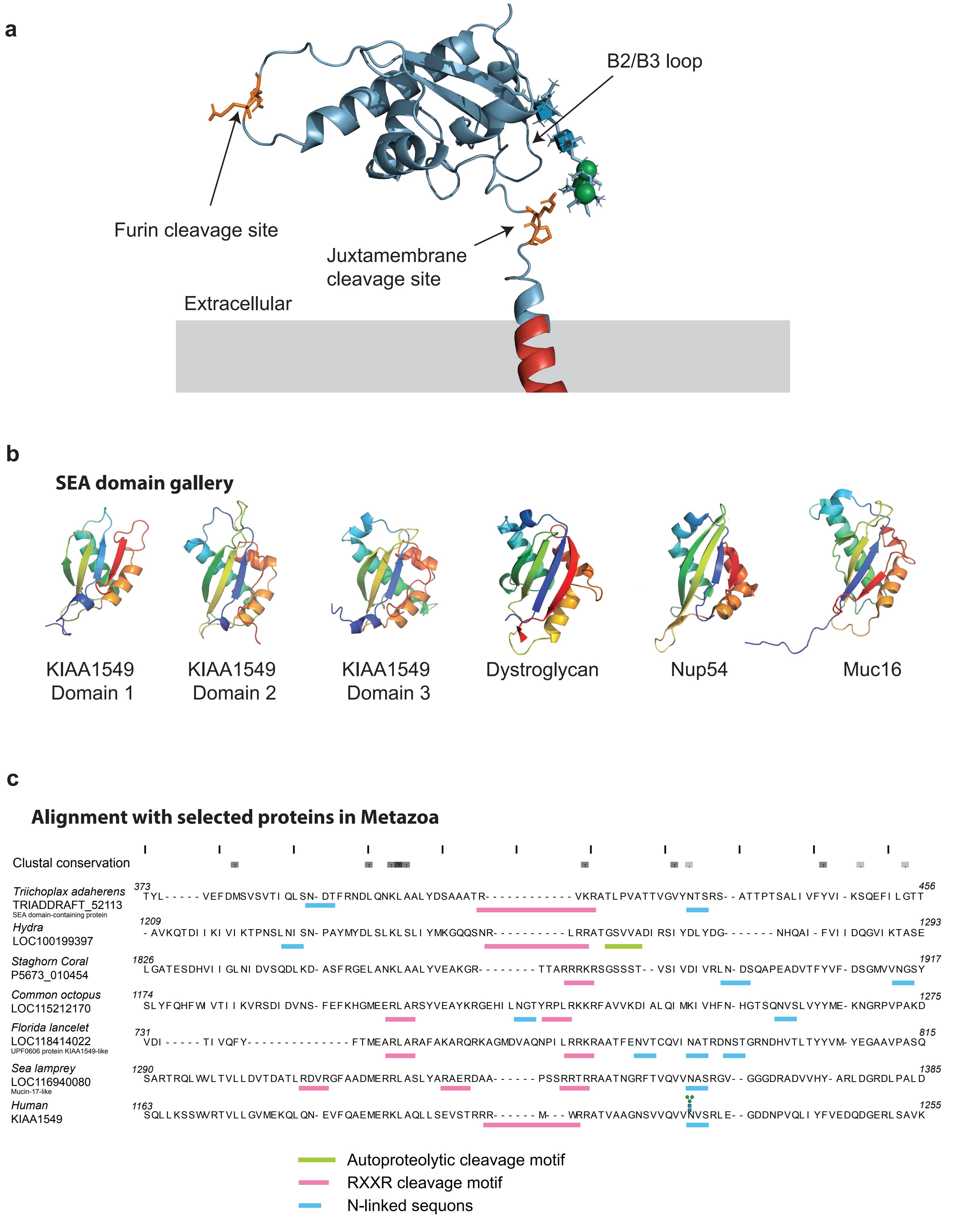

**Supplemental Figure 9. Pomt1 suppression reverses in vivo brain phenotype changes induced by K::B expression. (A)** Representative images of lower cortical regions of GFP control and K::B expressing samples. Note trapped neurons in the corpus callosum of K::B samples. **(B)** Additional images depicting changes in GFP+ transfected cell morphology within K::B + Rosa26 sgRNA and K::B + Pomt1 sgRNA samples. **(C)** INDEL percentage rate of CRISPR-Cas9 induced edits in Rosa26 or Pomt1 locus of neural stem cells isolated from Rosa26 sgRNA and Pomt1 sgRNA electroporated brains. Scale bars = 100um.

**Supplemental Figure 9**

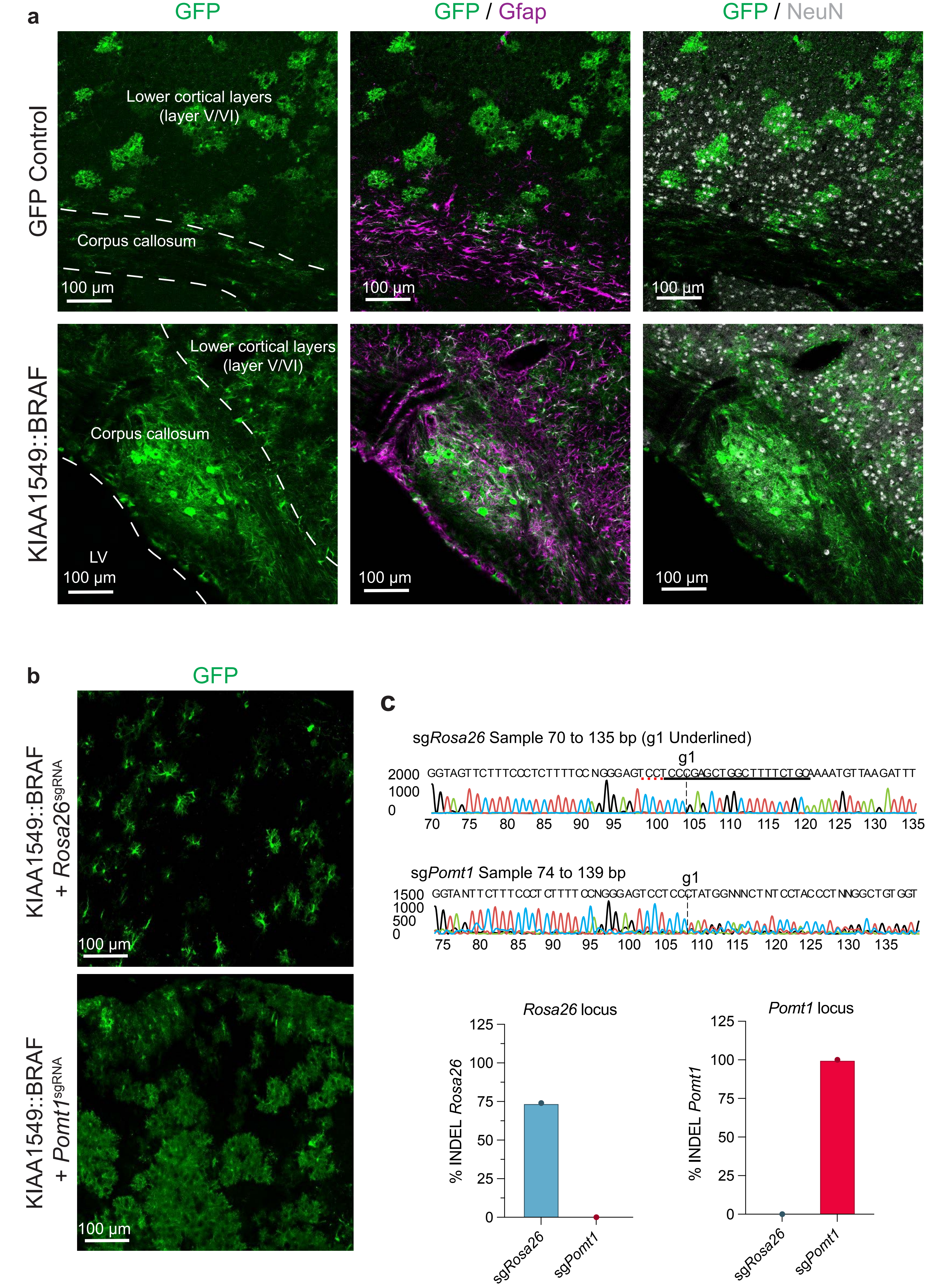
